# Directional purine–pyrimidine asymmetry in CRISPR repeats facilitates spacer acquisition

**DOI:** 10.64898/2026.07.20.739571

**Authors:** Sashikanta Barik, Parthasarathi Sahu, Koushik Ghosh, Hemachander Subramanian

**Affiliations:** Department of Physics, National Institute of Technology Durgapur, West Bengal, India 713209

**Keywords:** Purine–pyrimidine asymmetry, CRISPR–Cas systems, Spacer acquisition, Sequence-dependent asymmetric cooperativity, DNA unzipping kinetics

## Abstract

Spacer acquisition is the primary and essential step of CRISPR-Cas adaptive immunity in most prokaryotes and occurs preferentially at the leader-repeat junction of the CRISPR array. Despite the conservation of the adaptation machinery, spacer acquisition efficiencies and site-specificity vary markedly across CRISPR–Cas systems, suggesting that the local sequence architecture of the repeat and leader–repeat junction may contribute to the variation in the adaptation efficiency. To investigate this possibility, we systematically analyzed the direct repeat sequences and the terminal base pairs at the 3*^′^* end of leader regions adjacent to the integration site. We identify a conserved asymmetric purine-pyrimidine (RY) distribution within direct repeats, characterized by a 5*^′^* pyrimidine-rich half and a 3*^′^* purine-rich half, together with purine enrichment at the 3*^′^* end of leader sequences. Building on these observations, we develop a mechanistic model for spacer acquisition by demonstrating the importance of the sequence motif of the first direct repeat and the leader-repeat junction, using the sequence-dependent asymmetric cooperativity model that captures DNA unzipping kinetics. According to our model, sequences with such directional RY-asymmetric nucleotide distribution as direct repeats have high unzipping propensity and thus can promote efficient spacer integration. Consistent with this mechanism, experimental data from previous studies show that repeats with stronger asymmetry and appropriately positioned pyrimidine-to-purine transition sites are associated with larger spacer counts within CRISPR arrays. These findings reveal a conserved architectural feature of CRISPR repeats and provide a mechanistic model connecting DNA sequence organization to efficiency.

## 1 Introduction

The CRISPR–Cas system (Clustered Regularly Interspaced Short Palindromic Repeats and CRISPR-associated proteins) is an adaptive immune mechanism in most prokaryotes. It offers a sequence-specific defense against invading mobile genetic elements such as bacteriophages and plasmids [1, 2, 3, 4]. The core functional unit of this system is the CRISPR array, which consists of conserved direct repeats interspersed with unique spacer sequences, a leader sequence located upstream of the first direct repeat, and associated Cas genes. These spacers serve as molecular memory units of the previous infections and enable the system to recognize and neutralize future invasion by similar foreign genetic elements [5, 6, 7, 8, 9]. This defense mechanism operates through three distinct stages: adaptation, expression, and interference [10, 11, 12, 13]. During adaptation, short segments of the invader genome are captured and integrated as new spacers at the leader-proximal end of the CRISPR array by the Cas1–Cas2 adaptation complex, thereby encoding a heritable record of infection [14]. In the expression stage, the CRISPR array is transcribed into a precursor CRISPR RNA (pre-crRNA), which is then enzymatically cut and trimmed into individual, functional mature crRNAs. These crRNAs assemble with Cas proteins to form effector complexes that recognize and cleave complementary sequences in invading genetic elements during the interference stage [15].

Among the three stages of CRISPR–Cas immunity, the adaptation stage is the least understood, necessitating a detailed investigation into the mechanisms that control spacer integration into the CRISPR array. Spacer acquisition is a highly site-specific process, exhibiting a strong preference for the leader–first repeat junction, with only rare integration events occurring at other positions within or outside the array [16, 17]. The leader-repeat junction region contains regulatory elements crucial for the adaptation process, as demonstrated by the effect of site-specific mutations on spacer integration efficiency [18]. In addition, alterations within the first direct repeat itself significantly reduce integration efficiency, highlighting the importance of repeat sequence integrity in directing accurate spacer incorporation [19, 20].

Mechanistic insights into the site-specificity of spacer integration have emerged from structural, biochemical, and single-molecule studies, which collectively show that the process is mediated by the Cas1–Cas2 integrase complex [19, 21, 22, 23, 24, 25, 26]. Cas1–Cas2 selectively recognizes and binds prespacer DNA fragments derived from invading genetic elements, which often exhibit defined structural features such as 3*^′^* single-stranded overhangs. The integrase complex directs spacer insertion specifically at the leader-proximal end of the CRISPR array through a two-step transesterification reaction. In the first step, the 3*^′^* hydroxyl group of one prespacer strand nucleophilically attacks the phosphodiester backbone at the leader–repeat junction, resulting in strand integration. Subsequently, the second strand is incorporated at the repeat–spacer junction via a similar reaction, completing spacer insertion. Finally, host DNA repair pathways fill the resulting gaps and duplicate the flanking repeat sequence, thereby restoring array continuity and preserving CRISPR architecture [16, 18, 27].

The mechanism described does not provide clear insights into the site specificity of the adaptation process. Some studies indicate that the integration host factor (IHF) protein attaches to the A/T-rich leader sequence, directing spacer integration next to it [22, 21]. However, the tendency for new spacers to integrate at the leader-proximal repeat remains even without the integration host factor (IHF) [28, 29, 30, 18], which is generally absent in Gram-positive bacteria [31]. Consistent with this, integration events have been detected at the boundaries of multiple CRISPR repeats and in genomic regions near non-CRISPR palindromic sequences [32], indicating that intrinsic DNA sequence or structural characteristics may affect the choice of integration sites. Collectively, these findings suggest that the Cas1-Cas2 complex has an inherent capacity to identify specific sequence motifs or structural features of the CRISPR repeat and possibly the leader-repeat region [31, 33]. This inexplicability of site-specific integration, along with the potential role of the leader-repeat junction and the first direct repeat, calls for further research to uncover the underlying molecular mechanisms of the adaptation stage of CRISPR–Cas immunity.

In this study, we introduce a mechanistic framework to account for both the site specificity and the variability in spacer integration efficiency observed across CRISPR–Cas systems, even when similar adaptation modules (Cas1–Cas2 or Cas1–Cas2–3) are present. We analyze the CRISPR arrays, focusing on direct repeat sequences and the leader–repeat junction, to demonstrate how local sequence architecture affects the integration process. Our analysis uncovers a conserved asymmetric purine–pyrimidine nucleotide arrangement within direct repeats, characterized by a pyrimidine-rich 5*^′^* half and a purine-rich 3*^′^* half, along with purine enrichment at the 3*^′^* end of leader sequences near the integration site. Based on these observations, we propose a sequence-specific physical mechanism for spacer acquisition by utilizing the sequence-dependent asymmetric cooperativity model, which allows us to show how this intrinsic purine-pyrimidine asymmetry facilitates DNA unzipping [34, 35] and enhances efficient leader-proximal spacer integration.

## 2 Model

In this study, we utilize the *Asymmetric Cooperativity model* [36], whose central premise is the existence of an asymmetric influence of an existing base pair on the kinetic barrier heights of its right and left nearest-neighbor base pairs for their formation/dissociation. The asymmetric influence is of two kinds: the right asymmetric mode and the left asymmetric mode. In the right-asymmetric mode, the existing base pair lowers the kinetic barrier for formation and dissociation of its nearest neighbor base pair in the 3*^′^* direction (right), while simultaneously increasing the barrier for the adjacent base pair in the 5*^′^* direction (left). On the other hand, in the left-asymmetric mode, the kinetic bias is reversed, such that the pre-existing base pair reduces the kinetic barrier height of the adjacent base pair toward the 5*^′^* direction and raises the barrier of the base pair toward the 3*^′^*direction. Lower kinetic barriers facilitate faster base-pair formation/dissociation, whereas higher barriers hinder these processes. This right/left mode of a base pair depends on the *orientation* of the base pair, i.e., which base type, a purine (R) or a pyrimidine (Y), occupies which strand of the double-stranded DNA. Base pairs that are oriented as 3*^′^*-R-5*^′^*/5*^′^*-Y-3*^′^*catalyze their right nearest neighbors and inhibit their left nearest neighbors, where as, base pairs in 3*^′^*-Y-5*^′^*/5*^′^*-R-3*^′^* orientation catalyze their left nearest neighbors and inhibit their right nearest neighbors. The incorporation of asymmetric cooperativity requires base-pair complementarity and heteromolecular base pairing, and the cooperative influence is abrogated when both bases of a base pair are the same, due to symmetry considerations [36].

In contrast to the two modes of asymmetric cooperativity, dictated by the purine-pyrimidine orientation of base pairs in double-stranded DNA, single-stranded DNA also exhibits asymmetric cooperativity, which we have termed sequence-independent. This cooperativity is considerably stronger than the above sequence-dependent asymmetric cooperativity present in double-stranded DNA. The sequence-independent cooperativity operates during the construction of the daughter strand, where it catalyzes the formation of a new base pair in the 5*^′^* direction of the template, while inhibiting the dissociation of the already-formed base pair towards the 3*^′^*end. Such an arrangement of asymmetric cooperativity facilitates daughter strand construction and has therefore been argued to have been favored by evolution [37].

In the double-stranded DNA, the stronger sequence-independent influence is effectively canceled due to the antiparallel orientation of the two strands, resulting in zero net sequence-independent kinetic influence of a base pair over its neighbors. As a consequence, *sequence-dependent asymmetric cooperativity* emerges as the dominant mechanism governing the directional kinetic influence of a base pair on its neighbors in the double-stranded DNA [36].

## 3 Methods

In double-stranded DNA, *sequence-dependent asymmetric cooperativity* dictates the kinetic interactions between neighboring base pairs and thereby determines base-pair stability through modulation of kinetic barrier heights, as mentioned above. Base pairs associated with low kinetic barriers are more susceptible to dissociation and can act as initiation sites for strand unzipping, whereas base pairs with high kinetic barriers resist dissociation and impede the unzipping process and serve as termination sites of replication [34].

For example, in the sequence 3*^′^*-R*N_c_*Y-5*^′^*/5*^′^*-YNR-3*^′^*(where *N* denotes any nucleotide and *N_c_*its complementary base), both the left base pair 3*^′^*-R-5*^′^*/5*^′^*-Y-3*^′^* and the right base pair 3*^′^*-Y-5*^′^*/5*^′^*-R-3*^′^* lower the kinetic barrier height of the middle base pair 3*^′^*-*N_c_*-5*^′^*/5*^′^*-N-3*^′^*, making it the kinetically weakest site in the sequence. The middle base pair is therefore susceptible to dissociation and can trigger unzipping initiation, as demonstrated in Fig. 1a. In contrast, in the sequence 3*^′^*-Y*N_c_*R-5*^′^*/5*^′^*-RNY-3*^′^*, both the base pairs at both ends raise the kinetic barrier height of the central base pair, making it kinetically the strongest site, thereby suppressing the unzipping process (see Fig. 1a). As a result, the above sequence configuration slows strand-separation dynamics.

**Figure 1:**
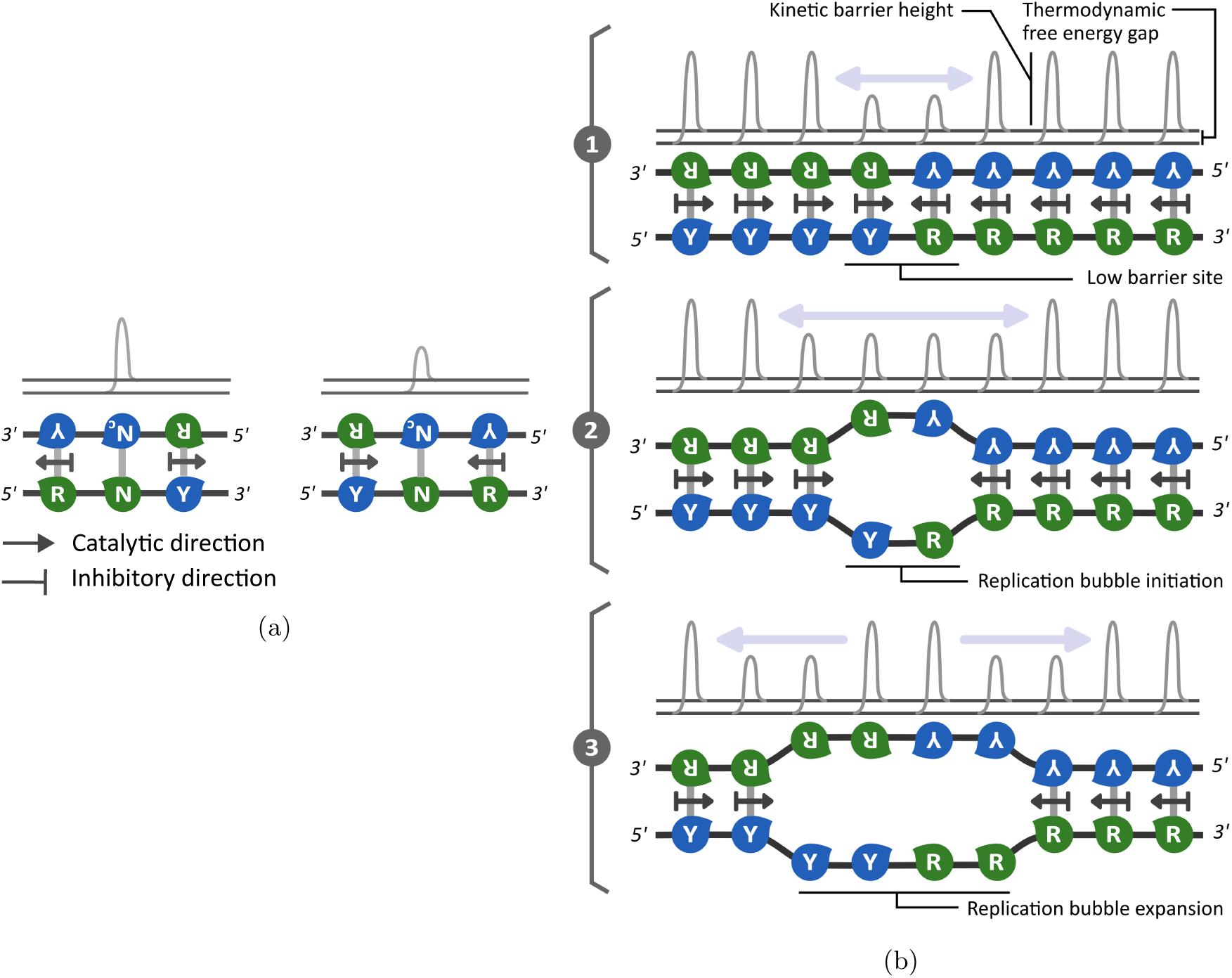
(a) Illustration of the asymmetric cooperative effect of a base pair on the kinetic barrier height of its two nearest neighboring base pairs. The asymmetric influence is indicated by an arrow mark for catalytic action and the bar head for inhibiting action. The base pair to the left, 3^′^-R-5^′^/5^′^-Y-3^′^ in the sequence 3^′^-RN_c_Y-5^′^/5^′^-YNR-3^′^ (N and N_c_are complementary base pairs), lower the kinetic barrier height of its right nearest-neighbor base pair (middle base pair), whereas, the base pair to the right, 3^′^-Y-5^′^/5^′^-R-3^′^, also lowers the kinetic barrier height of the central base pair between N and N_c_. The middle base pair, therefore, has the lowest kinetic barrier height and is prone to dissociation. However, the kinetic influence of the neighboring base pairs on the central base pair is entirely different in the sequence 3^′^-YN_c_R-5^′^/5^′^-RNY-3^′^. Both the base pairs at the edges increase the kinetic barrier height of the middle base pair, making it kinetically very stable, which therefore resists dissociation. The barrier heights are illustrated above the bonds for comparison. **(b)** Illustration of unzipping initiation in a high-skewed RY palindromic sequence, 5^′^-YYYYRRRRR-3^′^, of 9 bp length, based on the sequence-dependent asymmetric cooperativity model. The relative strengths of the base pairs are determined by their kinetic barrier height, indicated above each base pair. The base pairs with the lowest kinetic barriers are located at the center of the sequence and therefore initiate unzipping most readily (step 2). The variation of the barrier height of each base pair during the bidirectional unzipping propagation is shown in step 3.

**Figure 2:**
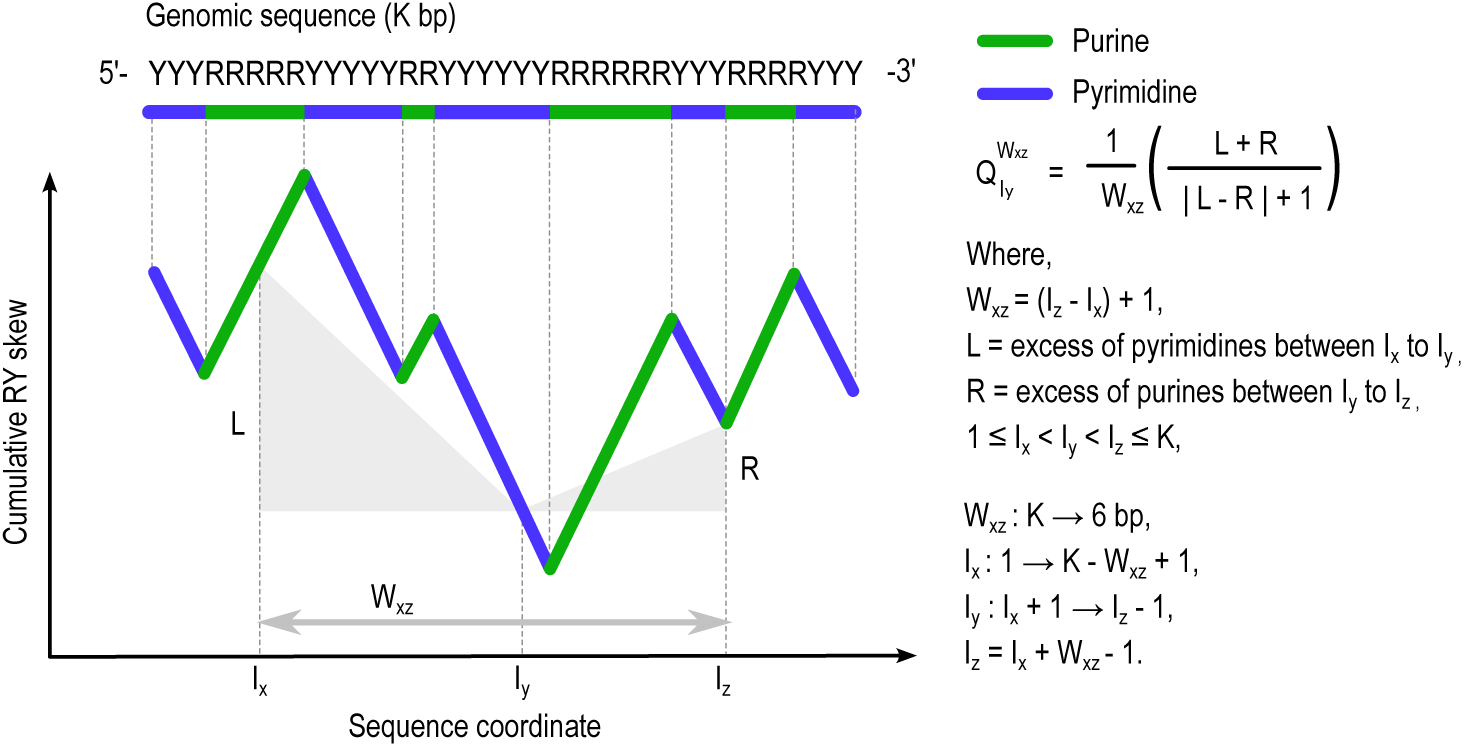
Schematic illustration of the cumulative RY skew profile of a genomic sequence for quantifying skew factor, the directional asymmetry of pyrimidine and purine distributions along 5^′^ → 3^′^. The genomic sequence is represented as a string of purines (R; A/G, green) and pyrimidines (Y; C/T, blue). Upward and downward trends in the cumulative RY skew profile indicate purine and pyrimidine enrichment, respectively. The skew factor has its maximum value for the sequence that displays strong compositional asymmetry (V - shaped skew profile), characterized by pyrimidine enrichment at the 5^′^ end and an equal-magnitude purine enrichment at the 3^′^ end. For the calculation of the skew factor, the sequence is scanned using three indices, I_x_, I_y_, and I_z_, satisfying 1 ≤ I_x_ < I_y_ < I_z_ ≤ K. A variable window, W_xz_ = I_z_ − I_x_ + 1, ranges from the full sequence length (K bp) to a minimum of 6 bp and, for each window size, is translated along the sequence in 1-bp increments. Within each window position, 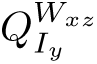 is evaluated at every intermediate index I_y_. The maximum value of 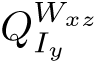 is retained for each window position and then normalized by the sequence lengt (K). The largest normalized value over all window sizes and window positions is defined as the skew factor.

According to the sequence-dependent asymmetric cooperativity model, the sequence 5*^′^*-YYYYRRRRR-3*^′^*(Fig. 1b) acts as an efficient unzipping initiation site (origin of replication) [34]. The base pairs in the left half of the sequence (3*^′^*-RRRR-5*^′^*/5*^′^*-YYYY-3*^′^*) operate in the right-asymmetric mode. They lower the kinetic barrier heights of their right nearest-neighbor base pairs (towards the 3*^′^* of the bottom strand) while raising the barriers to their left (towards 5*^′^*). On the other hand, the base pairs in the right half (3*^′^*-YYYYY-5*^′^*/5*^′^*-RRRRR-3*^′^*), which are simply 180*^◦^* rotated versions of the left half, operate in the left-asymmetric mode, exerting the opposite directional influence. As a result of these opposing cooperative biases, the central transition region becomes kinetically the weakest: the 4*^th^* and 5*^th^* base pairs (5*^′^*-YR-3*^′^*) have the lowest kinetic barrier heights, as illustrated in Fig. 1b. This occurs because the middle base pairs are catalyzed from both sides (e.g., in motifs of the form 5*^′^*-Y(Y)R-3*^′^*and 5*^′^*-Y(R)R-3*^′^*), making the central base pairs the preferred site for unzipping initiation, illustrated in step 2 of Fig. 1b.

Following the dissociation of the low-kinetic-barrier 4*^th^* base pair, the kinetic barrier of the 3*^rd^* base pair decreases due to the loss of inhibition from the now-non-existent 4*^th^* base pair, together with catalytic support from 2*^nd^* base pair, facilitating dissociation and promoting leftward (5*^′^* direction of the bottom strand) propagation of the unzipping fork. Similarly, the dissociation of the 5*^th^* base pair due to the catalyzing effect of the 6*^th^* base pair promotes rightward (3*^′^* direction) progression (step 3 of Fig. 1b). Consequently, regions governed by right-asymmetric cooperativity favor leftward fork movement, whereas regions governed by left-asymmetric cooperativity favor rightward fork movement. Together, these dynamics explain why sequences such as 5*^′^*-YYYYYRRRRR-3*^′^* exhibit high unzipping potential: strand separation initiates at the central pyrimidine-to-purine transition and propagates simultaneously in both directions, highlighting the functional role of the directional pyrimidine-purine asymmetric arrangement of nucleotides in promoting efficient bidirectional unzipping.

### 3.1 Modeling Unzipping dynamics

We utilize a continuous-time Markov chain (CTMC) model to quantify the association/dissociation dynamics of sequences such as those above. We define a Markov chain (*X_t_*)*_t≥_*_0_ with an associated state space *S*, whose elements are microstates representing distinct possible configurations of the double-stranded sequence, defined by different combinations of intact and disrupted base pairs of double-stranded DNA (dsDNA). The presence and absence of a base pair are denoted as 1 and 0, respectively, to construct the state space *S*. For example, a microstate for a dsDNA segment of length five base pairs with the rightmost base pair broken is represented as 11110. Under this binary representation of the state space, a dsDNA segment of length *n* base pairs possesses 2*^n^* (= M) possible configurational states, which can be used to characterize the DNA breathing process.

The dynamics of unzipping of a sequence is represented mathematically by the traversal of the Markov chain over the state space *S*. We impose the transition rule that breaking or forming a single base pair alone is allowed, i.e., only a single base pair can form or break. Following this transition rule, we construct a transition rate matrix Q with elements *q_ij_*, representing the transition rate from state *i* to state *j*, s.t. *i, j* ∈ *S*. The intrinsic rates of base-pair formation and dissociation in the absence of the nearest neighboring base pairs are denoted by *p* and *r* (thermodynamic rates), respectively, and when an adjacent base pair is present at either the 5*^′^* (left) or 3*^′^* (right) end, cooperative effects are incorporated by modulating *p* and *r* by multiplying appropriate kinetic parameters, *α* or *β*, depending on whether the neighbor inhibits or catalyzes the base pair under consideration. The catalytic effect of the neighboring base pair is taken into account through *α*, with *α >* 1, whereas the inhibitory effect is incorporated through *β*, with *β <* 1. When both the neighboring base pairs are present, the rates of base-pair formation and dissociation are modified multiplicatively by multiplying the bare formation/dissociation rates (*p* or *r*) by both *α* and *β*. The diagonal entries of the transition rate matrix are defined as 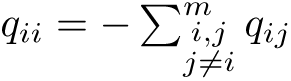 [38].

#### 3.1.1 Spectral decomposition analysis

Since our goal is to understand the unzipping dynamics of any given sequence, we need to extract the temporal unzipping behavior of the sequences from the transition rate matrix Q, which embeds all the necessary information needed for our purposes. We use the technique of spectral decomposition analysis to disentangle the dynamics of unzipping at multiple timescales, from the slowest to the fastest. Spectral decomposition involves eigen decomposition of the rate matrix Q to get eigenvalues and eigenvectors that carry information about the rates of different unzipping *modes* of the sequences under consideration. However, the rate matrix Q is not symmetric in our case, as the thermodynamic rates of base pair formation and dissociation are different, since the model represents a non-equilibrium situation. Diagonalization of this non-symmetric rate matrix will produce non-orthogonal left and right eigenvectors, and the eigenvalues of the matrix can be complex. We perform a symmetrization process through a well-known similarity transformation:

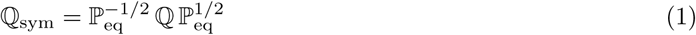

where P_eq_ is a diagonal matrix with the normalized left eigenvector **P**_eq_ corresponding to the eigenvalue zero of the asymmetric rate matrix Q along the diagonal; i.e., the eigenvector that describes the steady-state probability distribution of the Markov chain over all the states at infinite time, when the probability flow between all the states is balanced and detailed balance is satisfied. This can be verified from the equation

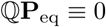

The eigenvalue equation for the symmetrized rate matrix is

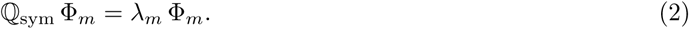

Here, *λ_m_*and Φ*_m_* represent the eigenvalues and orthonormal eigenvectors of Q_sym_. Since this non-equilibrium system eventually reaches steady state, the eigenvalues are all negative, except for the zero eigenvalue, which corresponds to the steady-state probability distribution at infinite time, with the eigen-vector **P**_eq_. These eigenvalues (*λ_m_*) are also the eigenvalues of the non-symmetric rate matrix Q. The left and right eigenvectors Ψ*_m_^L^* and Ψ*_m_^R^*, respectively, satisfy the following relation [39]:

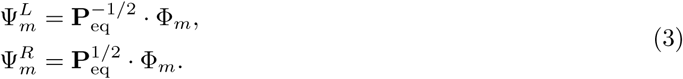

Here *m* represents the *m^th^* eigenvector or eigenmode of sequence unzipping, and each element of the eigenvector Ψ*_m_^L^*(or Ψ*_m_^R^*),Ψ*_m_^L^*(*i*), representing the *i^th^* element of the eigenvector, is related to the probability of occupation of the state *i* in the state space *S*, defined precisely below. The first right eigenvector, Ψ*_1_^R^*, corresponding to the eigenvalue *λ*_1_ = 0, provides the steady-state probability **P**_eq_, with the *i^th^* element of the eigenvector providing the probability of occupation of the state *i* at infinite time. Therefore, all the entries of this eigenvector are positive. The remaining eigenvectors contain both positive and negative entries. The sign of the eigenvector entries indicates the occupancy relative to the steady-state probability: positive entries are above **P**_eq_, whereas negative entries are below **P**_eq_ (Eq. 4) [40]. Therefore, the right eigenvector represents the probability distribution across the states of that dynamical mode, which grows or decays at the rate of the corresponding eigenvalue.

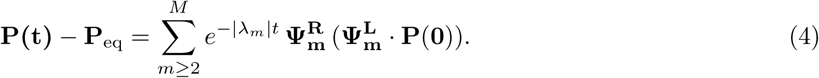

Here, **P**(**0**) is the probability distribution of the state occupancy of the Markov chain at *t* = 0, and **P**(**t**) provides the occupation probability of each state in the state space *S* (of size *M*) at time *t*. The left eigenvector is related to the right eigenvector by the transformation Ψ_m_^L=^ P_eq_ Ψ_m_^R^ [39], which is simply the projection of the right eigenvector onto the steady-state probability distribution. The *i^th^* element of the left eigenvector Ψ*_m_^L^* therefore represents how strongly the *i^th^* state contributes to the mode *m* when measuring from the steady state. For instance, the eigenvector Ψ_2_*^L^* of the smallest non-zero eigenvalue (*λ*_2_) provides insights into the contribution of each state to the state dynamics (breathing process) in the slowest mode of unzipping. To quantify how much each base pair in the sequence contributes to this slowest mode of unzipping, we define the following metric:

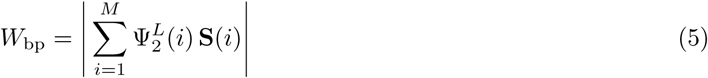

Here, **S**(*i*) is an element representing a state *i* in the state space *S*, that is, a vector of binary elements indicating the presence or absence of the hydrogen bond in a sequence. For example, **S**(*i*) = [1, 0, 1, 1, 1, 1] is a state of a six-length dsDNA sequence where the second hydrogen bond is broken. Since each element (state) of the left eigenvector Ψ_2_*^L^* measures the contribution of that element (state) to the slowest mode of unzipping of the dsDNA sequence, the above-defined sum, eq.5, provides us with a vector of length *n* representing the probability that a hydrogen bond in the sequence stays intact. This weight analysis of a dsDNA segment, therefore, provides a spatial profile of DNA breathing associated with the slowest timescale.

The values of the thermodynamic (*p* and *r*) and kinetic parameters (*α* and *β*) for this calculation are provided in the *Supplementary Information*.

### 3.2 Sequence analysis of the CRISPR array

To identify sequence motifs and characteristics responsible for spacer integration within the CRISPR array, we have performed sequence analysis of the direct repeats and the leader-repeat junctions. Two datasets were used: one comprising direct repeat sequences and another with base pairs at the leader–repeat junction. The direct repeat dataset, “direct repeat id.fsa”, was obtained from CRISPRCasdb [41, 42, 43] and contains 28,712 direct repeats with associated sequence-IDs but without corresponding leader sequences. To characterize nucleotide composition, direct repeats were centrally aligned, and the positional excess of purines over pyrimidines was computed at each sequence coordinate.

In parallel, we have also analyzed the leader–repeat junction sequences by extracting 10 base pairs from the 3*^′^*-end of the leader and 10 base pairs from the 5*^′^*-end of the first direct repeat. To construct this dataset, we sampled complete prokaryotic genomes from NCBI (https://www.ncbi.nlm.nih.gov/) and identified CRISPR arrays using the **CRISPRDetect** web server (http://crispr.otago.ac.nz/CRISPRDetect/ predict_crispr_array.html) [44]. The resulting dataset contains 500 CRISPR arrays annotated with accession numbers, repeat and spacer counts, along with their corresponding leader and repeat sequences.

#### 3.2.1 Cumulative RY Skew

To quantify the purine-pyrimidine distribution in a DNA sequence, nucleotides are grouped as purines (*A, G*) and pyrimidines (*C, T*). For a sequence *S_q_* of length *K*, we define a per-position score

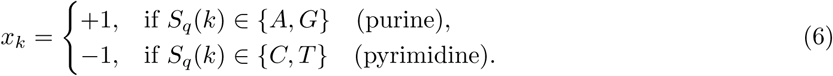

The cumulative RY skew is then computed as:

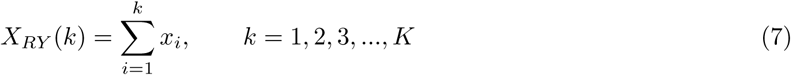

An increasing *X_RY_* (*k*) indicates local enrichment of purines, whereas a decreasing profile indicates enrichment of pyrimidines. When plotted along the sequence, this cumulative RY skew profile representation highlights the directional asymmetry in the nucleotide arrangement. For example, a sequence enriched in pyrimidines in the 5*^′^*-half and purines in the 3*^′^*-half produces a characteristic V-shaped profile with a minimum near the transition point Y→R, a signature frequently observed at replication origins of both prokaryotes and eukaryotes [34, 45].

#### 3.2.2 Skew factor

We further introduce another metric termed ‘skew factor’ to quantify the nucleotide symmetry of pyrimidine and purine distribution within each direct repeat along 5*^′^* → 3*^′^*. The metric is designed to become maximal when there is a large excess of pyrimidines in the 5*^′^* end and purines in the 3*^′^* end, and when these excesses are numerically equal. For example, it attains its maximum value for the sequence 5*^′^*-YYYYYRRRRR-3*^′^*, whereas it approaches zero for sequences that lack such a directional RY nucleotide arrangement, such as 5*^′^*-YYYYYYYYYY-3*^′^* and 5*^′^*-RRRRRYYYYY-3*^′^*. In addition, we define the *switching position* as the *relative position* along the 5*^′^* → 3*^′^* direction at which the nucleotide composition shifts from pyrimidine-to purine-enriched, scaled with respect to the sequence length. Following the sequence-dependent asymmetric cooperativity model, this *switching position* corresponds to a kinetically weak site with reduced barrier heights and is therefore expected to act as the initiation site for strand separation. For example, in the sequence 5*^′^*-YYYYYRRRRR-3*^′^*with the skew factor 1, the switching position lies at the 5*^th^* base pair, corresponding to a switching position of 5*/*10, from where unzipping initiates.

An algorithm (Fig. (S1)) was employed to compute the skew factor and identify the switching position, based on the following formula.

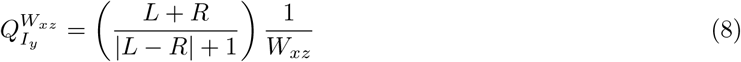

Here, 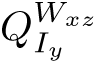 is calculated at an intermediate nucleotide position *I_y_* along the 5*^′^* → 3*^′^* direction within the window *W_xz_*.

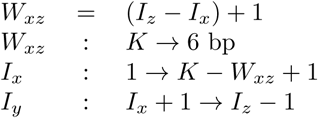

*L* quantifies the excess of pyrimidines over purines to the left (in the 5*^′^* direction) of *I_y_*, between *I_x_* and *I_y_*, and *R* quantifies the excess of purines over pyrimidines to the right (in the 3*^′^* direction) of *I_y_*, between *I_y_* and *I_z_*. Specifically, *L* = number of pyrimidines − number of purines between *I_x_^th^* and *I_y_^th^* nucleotides, and *R* = number of purines − number of pyrimidines between *I^th^* and *I^th^*nucleotides, within a window *W_xz_*.

The 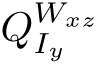 values are computed initially using a window *W_xz_* equal to the full sequence length, *K*, at each nucleotide position (*I_y_*). If the maximum value of 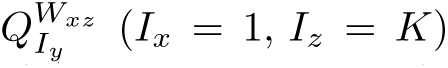 for this largest window size of K exceeds 0.9, i.e., *Q_max_* ≥ 0.9, then this value is taken as the skew factor, and its corresponding nucleotide position (*I_y_*) is identified as the switching position. If this criterion is not met, the window size is iteratively reduced by one nucleotide at a time down to a minimum length of 6 nucleotides. For each window size *W_xz_*, the window is translated along the sequence in single-nucleotide steps by varying *I_x_* from 1 to *K* − *W_xz_* + 1, with *I_z_* = *I_x_* + *W_xz_* − 1. Within each window position, 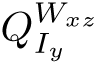 is evaluated at every intermediate nucleotide position (*I_y_*), where *I_x_ < I_y_ < I_z_*. The maximum value of 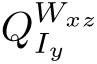 is then identified and normalized by the sequence length (*K*). This procedure is repeated for all window sizes and positions, and the largest normalized value is defined as the *skew factor*. The corresponding nucleotide position (*I_y_*) marks the switching position for the transition from the pyrimidine-rich to the purine-rich region of the sequence.

In general, the skew factor quantifies the similarity of a sequence to the directional RY-asymmetric palindromic motif (5*^′^*-pyrimidines-purines-3*^′^*) of the same sequence length and thus serves as an indicator for potential fast local unzipping sites. However, the skew factor also has certain limitations. For instance, it does not effectively distinguish between sequences with inherently low unzipping potential, and its discriminatory power decreases as sequence length increases due to the increase in degeneracy of skew factor values. These characteristics, specifically the relationship between skew factor and unzipping time, as well as the increasing degeneracy for sequences with low values of skew factor, are illustrated in Fig. S3.

## 4 Results

### 4.1 Asymmetric RY distribution in direct repeats

Sequence analysis of 28,712 direct repeats shows a pronounced RY-asymmetric nucleotide distribution, with a purine-rich 5*^′^* half and a pyrimidine-rich 3*^′^*half of the direct repeat, as illustrated in Fig. 3a.

**Figure 3:**
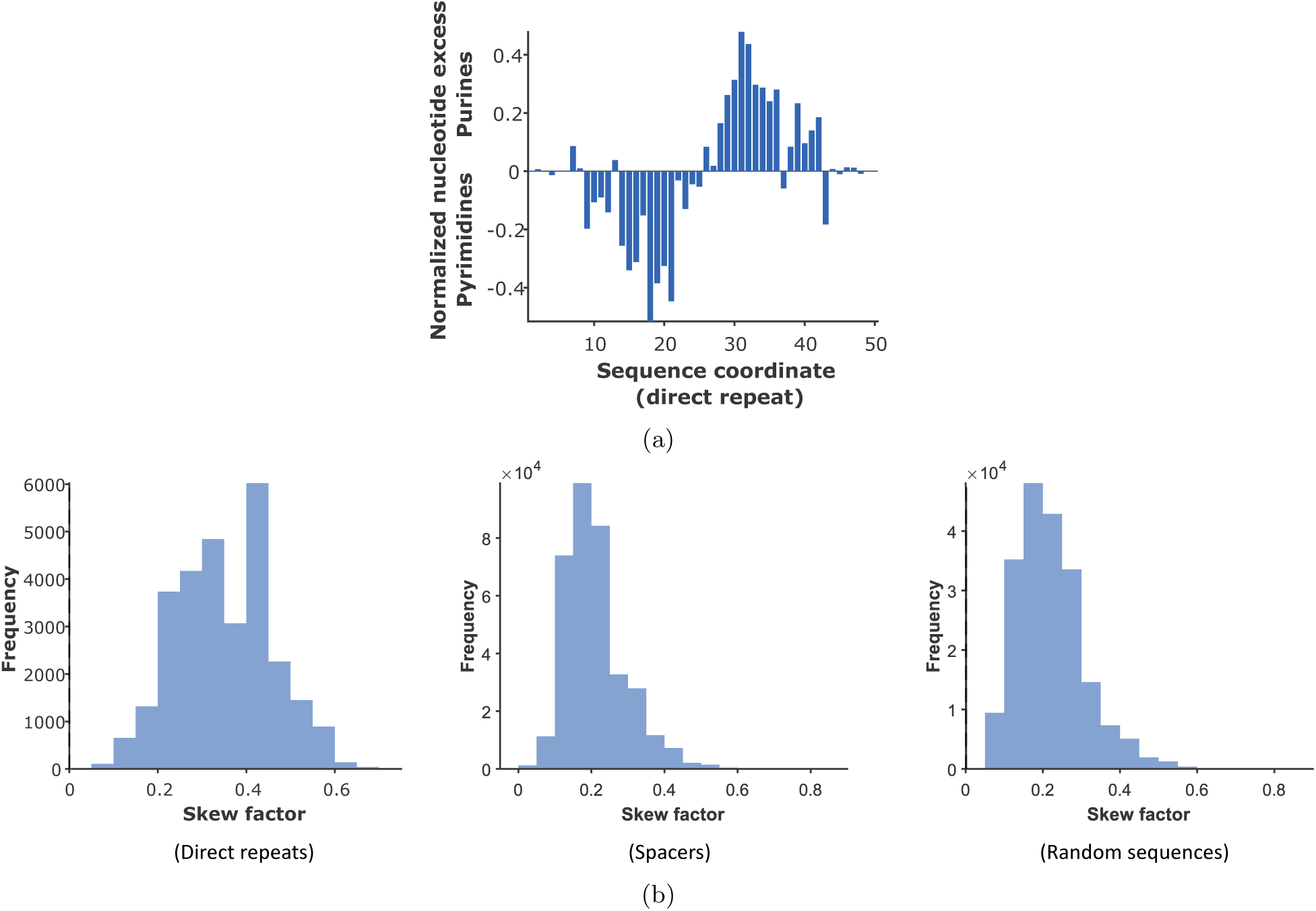
(a) The bar plot presents the normalized excess of purines over pyrimidines at each nucleotide position across 28,712 direct repeats. This sequence analysis was conducted by aligning the middle base pair of each direct repeat. A clear asymmetry is observed: the 5^′^ half of the sequence is enriched in pyrimidines, whereas the 3^′^ half is enriched in purines. This directional transition from pyrimidine-rich to purine-rich composition along 5^′^ → 3^′^ occurs near the midpoint of nearly every sequence. The uncertainty of the bar heights at both terminal ends is high, due to small number of repeats longer than 40 base pairs. **(b)** The skew factor distributions of CRISPR direct repeats (28,712 sequences) on the left, spacers (3,53,377 sequences) in the middle, and random sequences (2,00,000 sequences) on the right. The skew factor measures the directional RY asymmetry, the fraction of the sequence length exhibiting a directional transition from pyrimidines at the 5^′^ end to purines at the 3^′^ end (e.g., skew factor = 0.5 corresponds to 50% of the sequence length having 5^′^-pyrimidines-purines-3^′^ motif). The direct repeats have higher skew factors, with most values exceeding 0.2 and typically ranging between 0.2 and 0.6, indicating that directional RY asymmetry is commonly localized over ∼20–60% of the repeat length. While spacers and random sequences show substantially lower skew factors (mean ∼0.2), consistent with minimal or absent directional RY asymmetry.

Figure 3a presents a bar plot of the normalized positional excess of purines over pyrimidines across centrally aligned direct repeats. At each nucleotide position, the excess of purines over pyrimidines was computed across all contributing repeats and normalized by the number of sequences contributing at that position (sequence coordinate). This profile exhibits a pronounced directional purine-pyrimidine asymmetry, with the 5*^′^* half enriched in pyrimidines and the 3*^′^*half enriched in purines. A sharp transition near the center of the plot, at the sequence coordinate 25, marks the transition from a pyrimidine-rich to a purine-rich region.

To quantify the strength of this directional purine-pyrimidine asymmetry of each sequence, we have defined a metric called “skew factor” (see Methods) that measures the extent of 5*^′^*-pyrimidine to 3*^′^*-purine bias within each repeat sequence. For instance, a 10-base-pair sequence composed of five pyrimidines in the 5*^′^* half and five purines in the 3*^′^* half (5*^′^*-YYYYYRRRRR-3*^′^*) possesses a skew factor of 1. In contrast, a sequence arranged in the reverse orientation (5*^′^*-RRRRRYYYYY-3*^′^*) has a skew factor close to zero (0.08), reflecting asymmetry opposite to the defined directionality. The metric is designed in such a way that sequences lacking a clear positional enrichment of pyrimidines along the 5*^′^* direction and purines along the 3*^′^* direction exhibit skew factors approaching zero, indicating the absence of the defined RY asymmetry. For example, a skew factor of 0.2 indicates that 20% of the sequence displays such asymmetry. The presence of this nucleotide asymmetry is illustrated in Fig. 3b for direct repeats (28,712 sequences; left panel), spacers (353,377 sequences; middle panel), and random sequences (200,000 sequences; right panel). A majority of direct repeats exhibit skew factor values within the range 0.2–0.6. In contrast, both random sequences and spacers show markedly lower skew factor values, with averages near 0.2, indicating minimal or absent directional RY asymmetry.

### 4.2 Importance of asymmetric RY distribution in direct repeat for space integration

Further analysis of direct repeats shows an underlying relationship between the asymmetric RY distribution within the repeats and the number of spacers within the CRISPR array. The spacer count is found to depend on both the skew factor and the switching position of the pyrimidine-to-purine transition along the 5*^′^*–to–3*^′^* direction of the direct repeat. This relationship is illustrated in Fig. 4.

**Figure 4:**
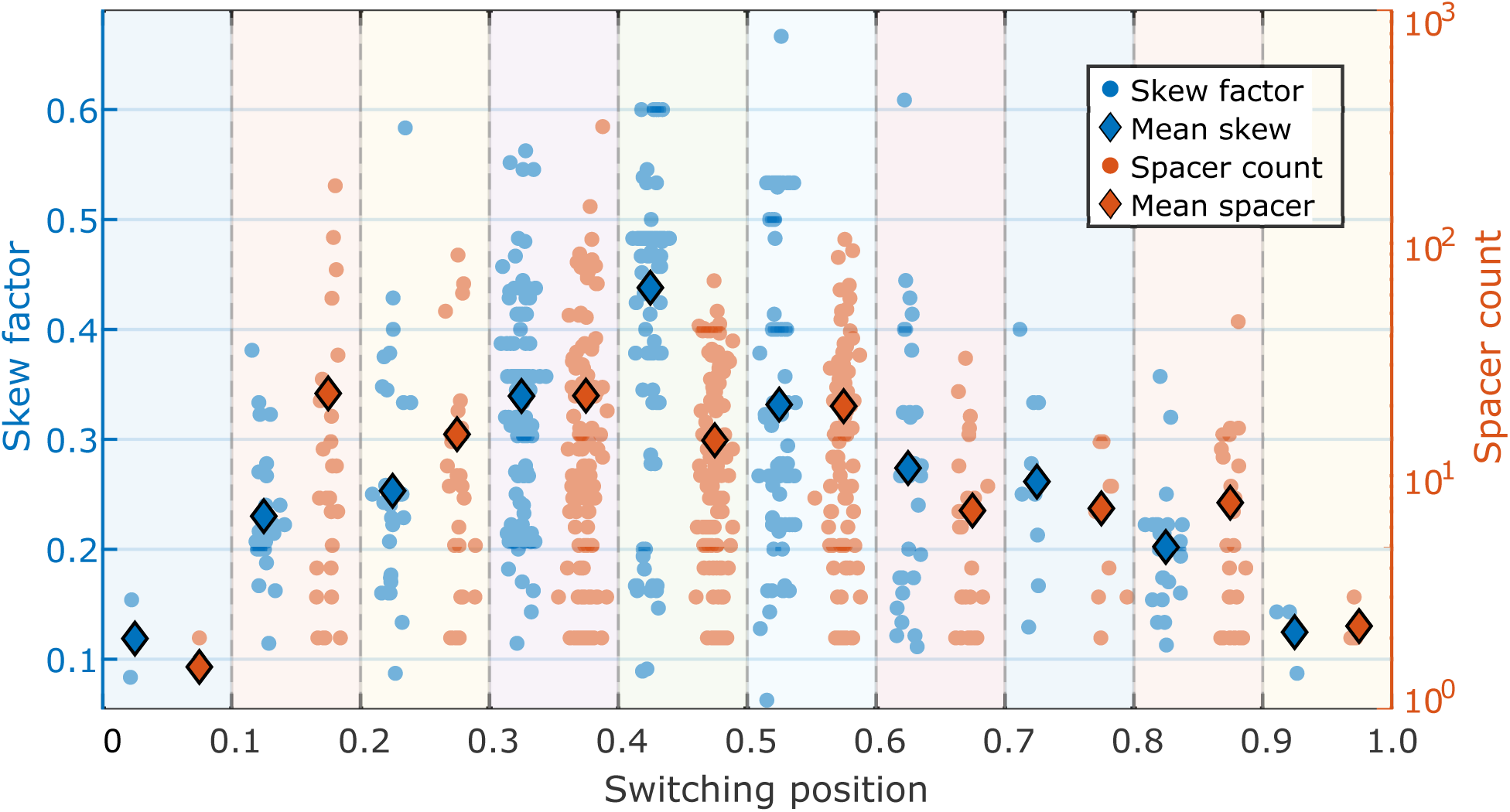
The figure illustrates the relationship between nucleotide directional RY asymmetry of a direct repeat and the number of spacers present in the CRISPR array carrying the direct repeat. The directional asymmetry is quantified by the skew factor, represented along the Y-axis with the blue dots, and the switching position range along the X-axis, marking the location for transition from pyrimidines to purines along the 5^′^ → 3^′^ direction. The corresponding spacer count is represented along the right Y-axis with the orange dots. The average values of the skew factor and the spacer count distribution within each switching position range are represented with blue and orange diamond markers, respectively. The figure shows that the direct repeats with a skew factor of more than 0.2 and switching position within 0.2-0.6 have a larger number of spacers than the others.

### Switching position

In Fig. 4, the x-axis denotes the switching position of the pyrimidine-to-purine (Y→R) transition, scaled by the corresponding direct repeat length. The left and right y-axes represent the distribution of skew factors and spacer count of the corresponding CRISPR arrays with blue and orange dots, respectively. The data show that CRISPR arrays with direct repeats exhibiting higher skew factors and switching positions located near the mid-region of the repeat tend to contain a larger number of spacers. Specifically, most direct repeats with switching position ranging from 0.2 to 0.6 have skew factors greater than 0.2 and are associated with substantially higher spacer counts. The direct repeats with switching positions near either end of the repeat, 0 to 0.2 toward the 5*^′^* end and 0.6 to 1 toward the 3*^′^* end, generally show lower skew factors and fewer spacers. Therefore, Fig. 4 suggests a functional relationship between the nucleotide motif of the direct repeat and the spacer acquisition efficiency.

### 4.3 Proposed mechanism of spacer acquisition

Motivated by the observed relationship between the nucleotide motif of the direct repeat and the spacer count, in Fig. 4, we aim to develop a generalized mechanistic framework to explain the spacer acquisition, based on the sequence architecture of the leader-repeat junction sequences and the direct repeats, in the light of the *sequence-dependent asymmetric cooperativity* model. This model helps us to understand the site-specific nature of spacer acquisition, using the sequence characteristics at the leader–repeat junction and the first direct repeat. The proposed mechanism (Fig. 5) illustrates the series of events from the unzipping initiation of the first direct repeat to repeat duplication during spacer integration.

**Figure 5:**
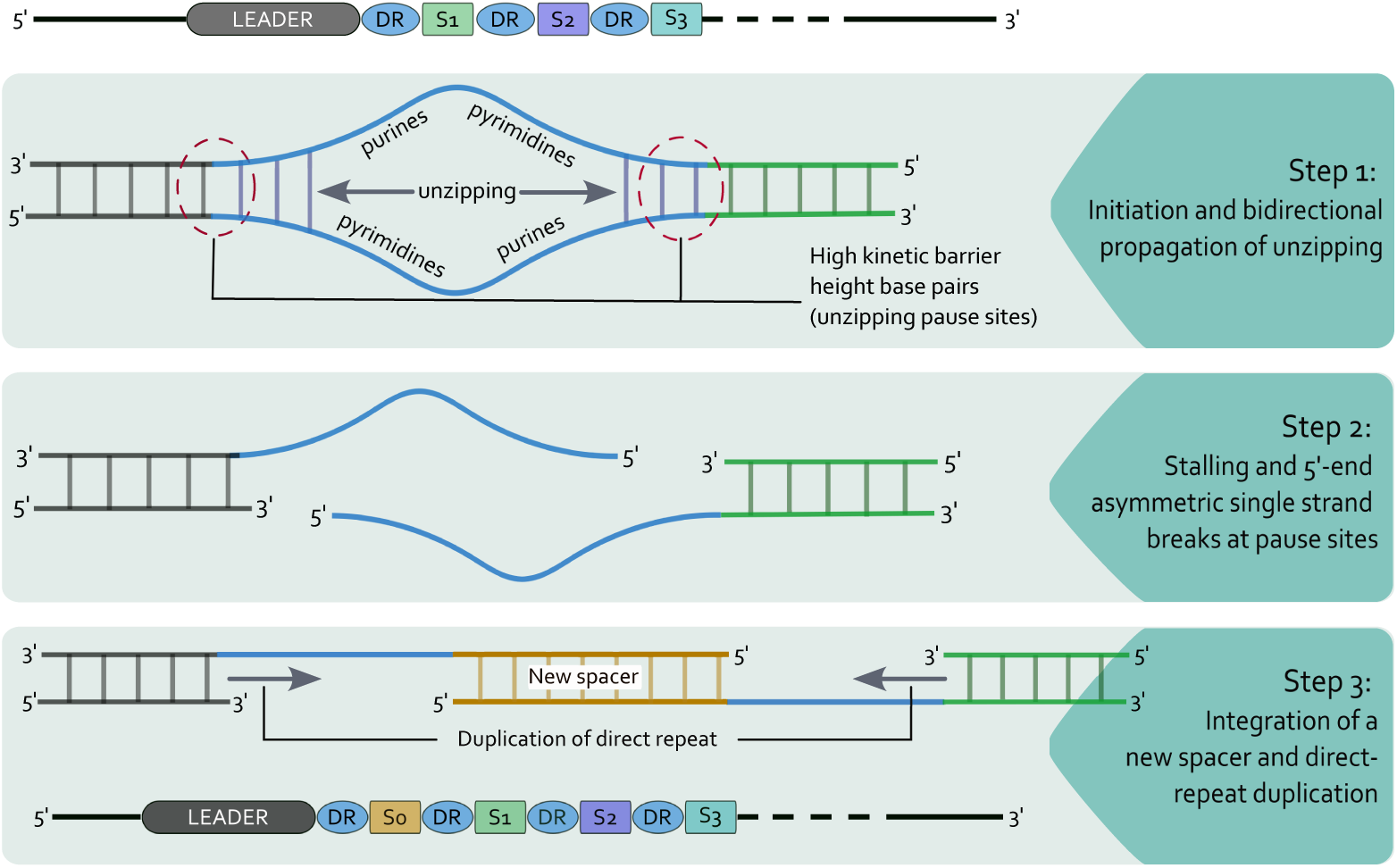
Illustration of our proposed mechanism underlying the spacer acquisition, which is based on the sequence analysis of direct repeats (in blue) and their leader sequences (in black) in light of the sequence-dependent asymmetric cooperativity model. Step 1: The RY-asymmetric nucleotide arrangement in the direct repeat, i.e., 5^′^-pyrimidines-purines-3^′^, is favorable for initiation and bi-directional propagation of the unzipping process. Step 2: Both ends of the first direct repeat have a high kinetic barrier, and hence impede the unzipping process, creating a replication bubble-like structure. This hybrid structure is intrinsically unstable, leading to asymmetric single-strand breaks at both 5^′^-ends of the direct repeat, thereby creating room for integration of a new spacer. Step 3: The new spacer gets integrated, and the spacer acquisition is completed with duplication of the direct repeats on both sides of the new spacer.

#### 4.3.1 Step 1: Initiation of unzipping in direct repeats

Figure 6a shows a V-shaped profile of the average cumulative RY skew computed over 28,712 direct repeats, obtained by aligning the sequences at their central nucleotide. The accompanying orange bars indicate the number of sequences contributing to each sequence coordinate. The 5*^′^* half of the direct repeat reflects a pyrimidine-rich region with an average excess of approximately 20% over purines, whereas the 3*^′^* half shows a purine-rich region with a comparable excess of approximately 20% over pyrimidines. This profile highlights a conserved nucleotide motif of the form 5*^′^*-*pyrimidines*–*purines*-3*^′^* within direct repeats.

**Figure 6:**
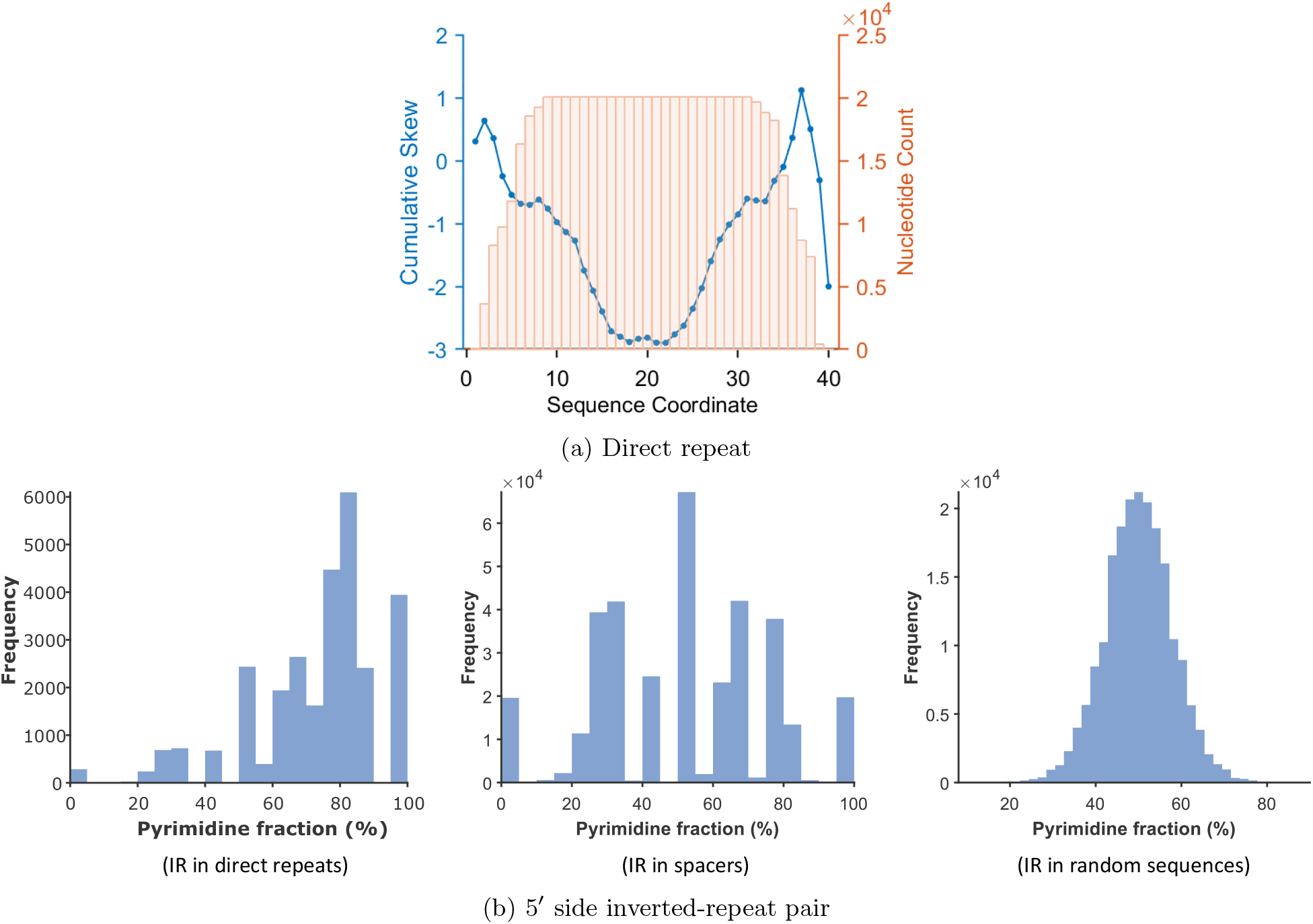
(a) Illustration of the average purine-pyrimidine cumulative skew profile of 28,712 direct repeats, aligned relative to their middle base pairs, while nucleotide count represents the number of sequences at each sequence coordinate via orange bars. The characteristic ‘V’ shape profile reflects an approximately 20% excess of pyrimidines in the left arm (5^′^ half) and an approximately 20% excess of purines in the right arm (3^′^ half). The vertex of the ‘V’ marks the switching position of the transition from pyrimidines to purines, where the base pairs are kinetically unstable, following the sequence-dependent asymmetric cooperativity model. An inverted ‘V’-shaped (Λ) local profile is observed at the 3^′^-end of the figure, indicating a transition from purines to pyrimidines. Base pairs at this site are kinetically resilient to dissociation. **(b)** The figure illustrates the distribution of pyrimidine fraction (%) of the 5^′^ arms of inverted-repeats, present within direct repeats (left), spacers (middle), and random sequences (right). 5^′^ arms of inverted repeats in direct repeats show a clear shift toward higher pyrimidine percentages, indicating consistent pyrimidine enrichment. In contrast, spacers and random sequences show an absence of nucleotide bias towards purines or pyrimidines.

The presence of this directional RY-asymmetric motif in a direct repeat can be attributed to the pyrimidine bias in the 5*^′^*arm of an inverted-repeat (5*^′^*-xxx-*R*_1_-xxx-*R*_2_-xxx-3*^′^*; *R*_1_ and *R*_2_ are reverse complements, representing the 5*^′^* and 3*^′^* arms, respectively.) within the direct repeat (Fig. 6b). Here, we analyzed the pyrimidine content of the 5*^′^* arm of the inverted repeat (arm length ≥ 3 bp) identified in each direct repeat, spacer, and random sequence. When multiple inverted repeats were present in a sequence, only the largest one was considered. Figure 6b shows the distribution of pyrimidine content in these 5*^′^* arms of inverted repeats identified in direct repeats (left), spacers (middle), and random sequences (right). The distribution for direct repeats is shifted toward higher pyrimidine content, indicating consistent pyrimidine enrichment in the 5*^′^* arm. In contrast, 5*^′^* arms in spacers and random sequences show no systematic pyrimidine bias. These results demonstrate that the presence of an inverted-repeat alone does not imply a pyrimidine-rich 5*^′^* arm (a purine-rich 3*^′^*arm).

Following the sequence-dependent asymmetric cooperativity model for dsDNA unzipping, we have shown that sequences with a 5*^′^*-*pyrimidines*–*purines*-3*^′^* distribution have high unzipping potential [34]. The unzipping profile for such sequences is illustrated in Fig. 1b. Ref. [34] observed that sequences with this directional RY-asymmetric motif exhibit faster unzipping rates than other sequences across a broad range of temperatures and sequence lengths, except at extremely low temperatures. The unzipping times of the fastest and slowest sequences for different lengths and temperatures are shown in Fig. S2. These findings suggest that sequences possessing 5*^′^*-pyrimidines–purines-3*^′^* motifs are intrinsically capable of participating in unzipping initiation and are therefore well suited to function as direct repeats during spacer integration into CRISPR arrays.

#### 4.3.2 Step 2: Stalling of the unzipping process

Unzipping in the sequences with the directional RY asymmetric distribution of nucleotides (5*^′^*-pyrimidines-purines-3*^′^*) initiates from the central base pairs and propagates in both directions simultaneously (see Fig. 1b). This bidirectional unzipping of the direct repeat is impeded at the base pairs with high kinetic barrier heights. The base pairs at the transition point of the nucleotide composition from purines to pyrimidines along the 5*^′^*→ 3*^′^* direction exhibit the maximum barrier heights. For example, in sequence motifs such as 5*^′^*-RNY-3*^′^* (Fig. 1a), the middle base pair has the highest kinetic barrier height because both neighboring base pairs, purine in the 5*^′^* end and pyrimidine in the 3*^′^* end, are inhibiting its dissociation. Such purine-to-pyrimidine transitions are observed near the 3*^′^* end of direct repeats, as illustrated in Fig. 6a, where a Λ-shaped local skew profile appears toward the right side of the cumulative RY skew profile of the direct repeat near sequence coordinate 37. Pyrimidines, specifically cytosines, at the 3*^′^*-end of the direct repeat, as a repeat triplet, are also observed [46, 47].

In addition to the 3*^′^* end of the direct repeat, a purine-to-pyrimidine transition is also observed at the leader-repeat junction, as shown in Fig. 7. This figure shows the average cumulative RY skew profile of the leader-repeat junction sequences, which are obtained by concatenating the terminal 35 base pairs from the 3*^′^*end of 500 leader sequences (in blue) with the initial 15 base pairs from the 5*^′^*end of their corresponding first direct repeats (in orange). The upward trend in the skew profile represents the excess of purines, while the downward trend corresponds to the pyrimidine excess. The leader-side skew profile exhibits an upward trend, while the repeat-side skew profile exhibits a downward trend. A local Λ-shaped skew profile is present near the leader-repeat junction region (at sequence coordinate 36), which marks the transition of purines to pyrimidines. This transition corresponds to the location of base pairs with high kinetic barrier heights. Taken together, Fig. 6a and 7 demonstrate the presence of high–kinetic-barrier base pairs at both ends of the direct repeat, which impede the unzipping.

**Figure 7:**
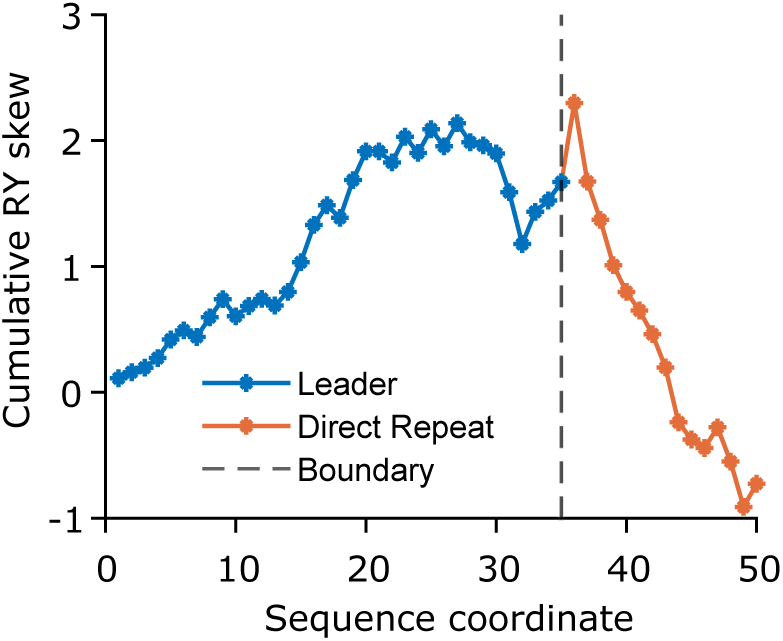
The figure shows the average cumulative RY skew profile of 500 leader-repeat junction sequences, assembled by concatenating 35 base pairs from the 3^′^-end of the leader sequence (in blue), and 15 base pairs from the 5^′^-end of the first direct repeat (in orange). The Λ shape skew profile near the junction (dashed vertical line), at sequence coordinate 36 (first base pair of direct repeat), shows the transition of purines to pyrimidines, indicating the presence of high-barrier-height base pairs at the leader-repeat junction. The leader-side skew profile shows an increasing trend, indicating an excess of purines over pyrimidines.

We perform a quantitative analysis to demonstrate that the presence of a Λ-shaped skew profile in the cumulative skew diagrams of leader-repeat junctions and direct repeats results in inhibition of the unzipping process. In the CTMC simulation of the unzipping of dsDNA, we performed the spectral decomposition analysis of the unzipping process (see the *Methods*). The persistence or stability of each base pair in the slowest mode is calculated using Eq. 5 and illustrated in Fig. 8 for two sequences. In the sequence 5*^′^* − *RRYYYRRRY Y* − 3*^′^*, which has similar skew profile as that of the direct repeat, we observe that 2*^nd^,* 3*^rd^,* 8*^th^* and 9*^th^* base pair contribute the most to the unzipping process. Similarly, in the sequence 5*^′^* − *YYRRRYYYRR* − 3*^′^*, the middle base pairs (5*^th^* and 6*^th^*) make the largest contribution. Consequently, these base pairs persist longer than others during the unzipping and can delay the strand separation.

**Figure 8:**
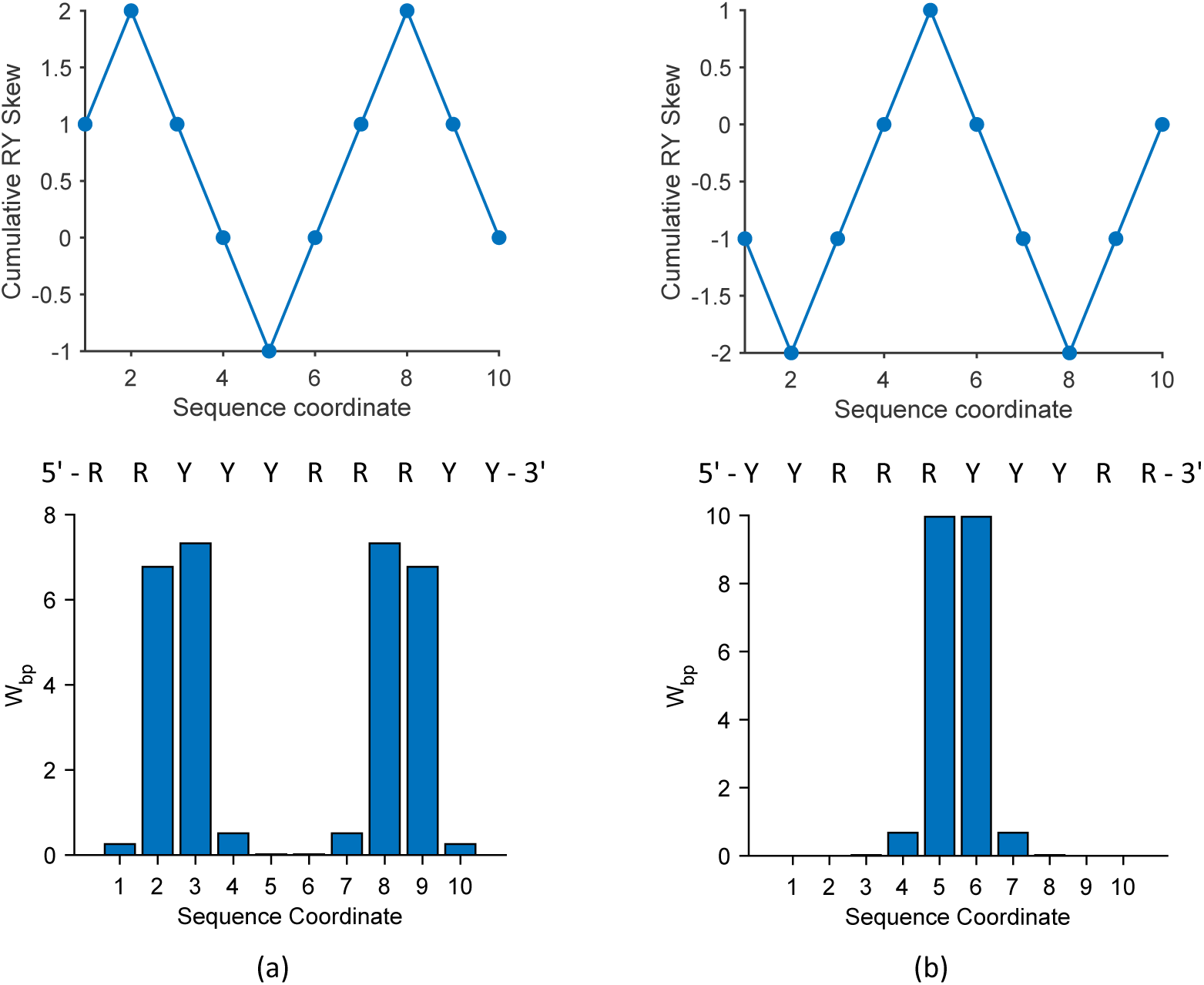
The unzipping dynamics, at the slowest mode, for two representative dsDNA sequences: (a) 5^′^–RRYYYRRRYY–3^′^ and (b) 5^′^–YYRRRYYYRR–3^′^. These sequences have motifs similar to those present in the sequences analyzed throughout the manuscript. In each panel, (a) and (b), the upper plot shows the cumulative RY-skew profile of the sequence, and the lower plot illustrates the contribution of each base pair to the unzipping process in the slowest mode. The weights of each base pair (W_bp_) during the unzipping process, which measures the difficulty of breaking of the base pairs, are shown on the Y-axis, with the corresponding base-pair positions on the X-axis. The base pairs positioned at the Λ-shaped skew profile are more difficult to break because their dissociation kinetic barrier height is high, and they persist longer during unzipping.

#### 4.3.3 Step 3: Integration of a new spacer

The direct repeat forms a bubble, a hybrid structure of dsDNA:ssDNA, due to the stalling of the unzipping process, caused by the kinetic bottleneck at both ends of the first direct repeat, as demonstrated above. Such a configuration of DNA is unstable and prone to single-strand breaks [48, 49, 50]. The single-strand breaks occur at both 5*^′^*-ends of the bubble near the junction of dsDNA and ssDNA, where the 5*^′^*-ends of the single strands inside the bubble are pyrimidine-rich and the 3*^′^*-ends are purine-rich (Fig. 5). Such asymmetric single-strand breaks, 5*^′^*-end resection, are observed during genomic activity like recombination/rearrangement [51, 52, 53]. Furthermore, Cas1 protein from Escherichia coli (YgbT) has been reported to show nuclease activity toward single-stranded and branched DNA substrates, including Holliday junctions, replication forks, and 5*^′^*-flaps [54]. The single-strand breaks at both 5*^′^*ends of the first direct repeat in the CRISPR array create room for the new spacer to integrate, as illustrated in Fig. (5).

Alternatively, the transient bubble-breathing dynamics of this hybrid intermediate may promote recognition by the adaptation machinery (Cas1–Cas2). Cas1–Cas2 can then bind and orient the two 3*^′^*-OH ends of the prespacer as nucleophiles, which sequentially attack the target phosphodiester backbone at the leader–repeat and repeat–spacer junctions, driving two transesterification reactions that covalently integrate the prespacer. Finally, host repair enzymes fill the resulting gaps, seal nicks, and duplicate the adjacent repeat to restore CRISPR array integrity.

## 5 Experimental Support

The proposed mechanism emphasizes that sequence-dependent unzipping kinetics is the key determinant of direct repeat functionality during spacer integration. In particular, direct repeats exhibiting a 5*^′^*-pyrimidine–purine-3*^′^* nucleotide arrangement possess high unzipping potential, enabling efficient initiation and simultaneous bidirectional propagation of strand separation within the repeat. While purine-to-pyrimidine transitions, 5*^′^*-purine–pyrimidine-3*^′^*motifs, present near both ends of the first direct repeat (the leader–repeat and repeat–spacer junctions), can kinetically hinder further strand separation. Additional sequence motifs can play analogous roles by either promoting or stalling unzipping dynamics. A comprehensive list of high-and low-unzipping potential motifs, obtained from mean first passage time (MFPT) analysis of the sequences using a continuous time Markov chain as described in the Methods section, is provided in Fig. S2.

Sequences containing one or more pyrimidine-to-purine transition stretches along the 5*^′^* → 3*^′^* direction (e.g., 5*^′^*-YYYYYRRRRR-3*^′^*, 5*^′^*-YYYYRYRRRR-3*^′^*, and 5*^′^*-YYYRRRYYRR-3*^′^*) exhibit a high unzipping propensity (Fig. S2). In contrast, alternating purine–pyrimidine arrangements (e.g., 5*^′^*-RYYRYYRYYR-3*^′^*, 5*^′^*-RYRYRYRYRY-3*^′^*, and 5*^′^*-RRYYRRYRYY-3*^′^*) substantially reduce the unzipping rate, whereas purine-to-pyrimidine transitions, such as 5*^′^*-RRRRRYYYYY-3*^′^*, can effectively stall the unzipping process owing to a reversal in the mode of cooperativity.

For example, in the sequence 5*^′^*-RRYYYRRRYY-3*^′^*(Fig. 8a), unzipping initiates at the central base pair and propagates bidirectionally. This behavior arises from the right-asymmetric mode of cooperativity of the pyrimidine tract on the 5*^′^* side and the left-asymmetric mode of cooperativity of the purine tract on the 3*^′^* side (*Methods*). However, the propagation is impeded by changes in the cooperativity mode at the 2*^nd^* nucleotide from the 5*^′^* end, where the cooperativity becomes left-asymmetric, and at the 8*^th^* nucleotide, where it switches to right-asymmetric on the 3*^′^* side. These reversals disrupt the directional propagation of the unzipping front, thereby hindering further unzipping.

This study of unzipping dynamics helps us to reinterpret the experimental observations of the site-specific nature and efficiency of spacer acquisition by a CRISPR array. By examining the effects of reported mutations in the CRISPR array through this kinetic lens and using representative model motifs (see Fig. S2 for the fastest and the slowest sequence motifs), we provide experimental support for our proposed mechanism underlying both acquisition efficiency and integration site specificity.

### 5.1 Mutational effect on the site-specific nature and the efficiency of spacer acquisition

A broad range of experimental studies consistently identifies the leader–repeat junction as the dominant integration site for new spacers, highlighting the functional importance of the leader-repeat junction in directing site-specific acquisition [23, 55, 56, 14]. The observations in Refs. [20, 18] suggest that a conserved segment of approximately 10 base pairs at the leader–repeat junction is essential for successful spacer integration, which aligns with the predictions of our kinetic model. Following our model, the leader–repeat junction behaves as a checkpoint for unzipping because of its nucleotide arrangement (see Fig. 7). Such stalling of local unzipping could facilitate the insertion of new spacers (Fig. 5).

#### 5.1.1 Sequence preferences at the leader–repeat junction

Kim *et al.* [20] analyzed the nucleotide preferences at the leader–repeat junction during spacer integration using two plasmid variants. One variant had the leader and repeat–spacer units from the *S. thermophilus* CRISPR array (pCRISPR), and the other plasmid did not contain the CRISPR array (pControl). In both plasmid variants, integration sites exhibited a strong purine bias, near the 3*^′^*-end of the leader. In pCRISPR, guanine (G) and adenine (A) were preferred at 56.6% and 22.1%, respectively, whereas cytosine (C) and thymine (T) accounted for only 13.7% and 7.7%. A similar distribution was observed in pControl (G: 54.1%, A: 21.1%, C: 17.1%, T: 7.7%), indicating a preference for purines in the local integration sequence context.

We further analyze the leader-repeat junction sequence 5*^′^*-CATTTGAG-GTTTTTGTA-3*^′^* (leader: 5*^′^*-CATTTGAG-3*^′^*; repeat: 5*^′^*-GTTTTTGTA-3*^′^*) used in Ref.[18]. Spacer integration occurred near the center of this sequence (9*^th^* and 10*^th^* base pairs) at the leader–repeat boundary. This sequence composition reflects a purine-to-pyrimidine transition at the junction, consistent with our predicted unzipping pause site associated with high kinetic barriers (Fig. 7). Mutations in this region, changing 5*^′^*-CATTTGAG-GTTTTTGTA-3*^′^* to 5*^′^*-AAGTCATGC-GTTTTTGTA-3*^′^* or 5*^′^*-ATGCCATTT-GTTTTTGTA-3*^′^*, abolished the spacer integration at the junction. These mutations disrupt the purine enrichment in the leader segment, which is required to maintain the high kinetic barrier heights at the leader–repeat junction, reinforcing our model prediction that spacer integration may be influenced by the local nucleotide arrangement and its impact on unzipping kinetics, rather than solely by the presence of a CRISPR array.

#### 5.1.2 Sequence motifs near spacer integration hotspots

To further validate our proposed mechanism underlying the site specificity and the efficiency of the spacer acquisition, we examine the experimentally identified integration sites with corresponding reads of the insertion events within the CRISPR arrays, which are composed of a few 3*^′^*-end base pairs of the leader (blue dots), first direct repeat (red dots), and a portion of the first spacer (green dots), as illustrated in Fig. 9. For this analysis, we used experimental datasets from Grainy *et al.* [29] and Kim *et al.* [20], shown in Figs. 9a and 9b, respectively. These figures show the cumulative RY-skew profiles of the CRISPR arrays, with each panel annotated with the reported mutation type (as used in those studies). The bar plots in these figures (right Y-axis) represent the reads in percentage, where the black bars are for the leader-repeat insertion sites and the gray bars are for the repeat-spacer insertion sites.

**Figure 9:**
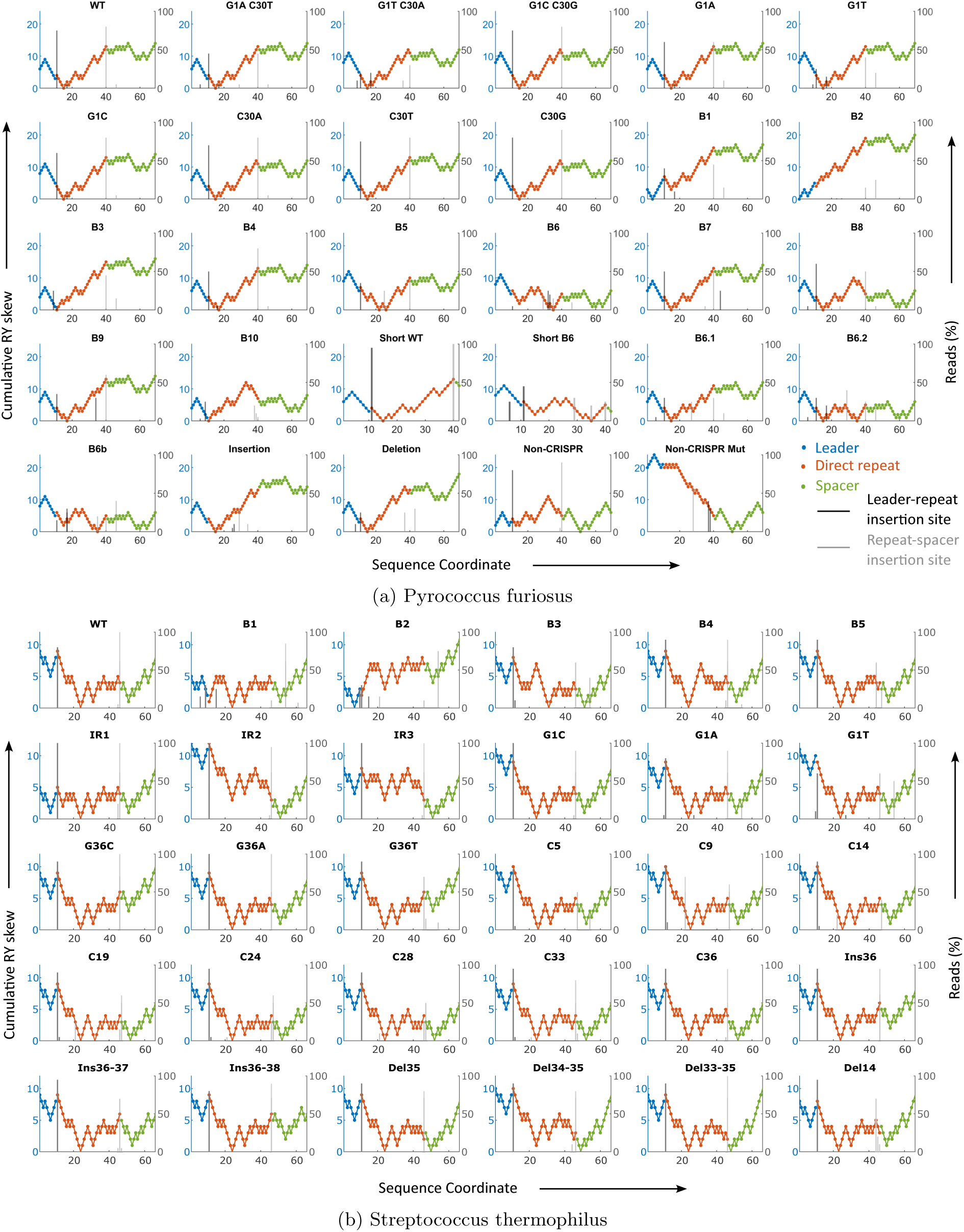
The figure illustrates the cumulative RY skew profiles (left Y-axis) of the CRISPR arrays from two experimental studies, together with the insertion sites of newly integrated spacers and the corresponding read counts of the insertion events in a bar plot (right Y-axis). The CRISPR array is composed of the leader sequence (blue dot), the first direct repeat (orange dot), and the first spacer sequence (green dot). The position of the bars marks the insertion sites. The dark bars represent the primary insertion sites (leader-repeat insertion sites), while the gray ones represent the secondary insertion sites (repeat-spacer insertion sites). The height of the bars indicates the percentage of the read count. The experimental data used in (a) is from [29] and (b) is from [20].These plots illustrate that the presence of Λ-shaped skew profiles are necessary for a site to function as an insertion site, but not sufficient.

Across both experimental datasets (Figs. 9a and 9b), the cumulative RY-skew profiles of most CRISPR arrays exhibit one or two pronounced V shapes, corresponding to pyrimidine-to-purine transitions along the 5*^′^* → 3*^′^* direction. In contrast, the arrays lacking these characteristic features tend to exhibit weakened or poorly defined integration hotspots, as indicated by short bars. Following our proposed mechanism, insertion events preferentially localize near regions associated with high kinetic barrier heights, such as Λ-shaped skew profiles due to purine-to-pyrimidine transitions and zig-zag patterns (5*^′^*-RYRYRYRYRY-3*^′^*) due to multiple RY transitions in the cumulative RY skew profile. The integration sites shift with the positional rearrangement of these low-and high-barrier regions, indicating that the distribution of kinetically favorable and unfavorable sites for unzipping within the leader–repeat–spacer segment can modulate both the efficiency and location of spacer integration. A detailed analysis of Fig. 9 is included in the *Supplementary Information*.

The observations in Fig. 9b support our proposed mechanism, and illustrate the fact that the Λ-shaped skew profiles are necessary, but not sufficient for a position to function as an insertion site. However, Fig. 9a shows some discrepancies in explaining both the site-specificity and the efficiency of the spacer integration. One possible explanation for these discrepancies is that the experiments in Ref. [29] were performed using the hyperthermophilic archaeon Pyrococcus furiosus. The elevated temperatures associated with hyperthermophilic systems can facilitate the dissociation of thermodynamically weak base pairs, thereby altering DNA unzipping dynamics and potentially affecting spacer integration. In addition, previous studies have shown that naive adaptation in Pyrococcus furiosus occurs with relatively low basal efficiency, whereas Streptococcus thermophilus exhibits comparatively robust spacer acquisition activity [57].

#### 5.1.3 Influence of nucleotide arrangement on adaptation efficiency

We next analyze the dependence of adaptation efficiency on the nucleotide arrangement of the CRISPR array, using the data reported in Refs. [27, 58]. In the reported data, the CRISPR arrays were generated by substitution mutations of the sequence 5*^′^*-GTGT-TCCCCGCGCCAGCGGGGA-TAAACC-3*^′^* (5*^′^* −*Leader* − *Direct Repeat* − *Spacer* − 3*^′^*), for which adaptation efficiencies vary across mutants [58]. We examined CRISPR array mutants by comparing the nucleotide motifs of the arrays with experimentally measured adaptation efficiencies.

Figure 10a reveals a positive correlation between the skew factor of the direct repeat and log(adaptation efficiency), suggesting that arrays with lower skew factors have reduced adaptation efficiency, while those with higher skew factors exhibit enhanced adaptation efficiency. This strong correlation likely results from the presence of two distinct clusters. The diagonal arrangement of these clusters also suggests the same notion that higher skew factors may be necessary for high adaptation efficiency. Notably, a similar relationship between the skew factor and adaptation efficiency is observed within the cluster.

**Figure 10:**
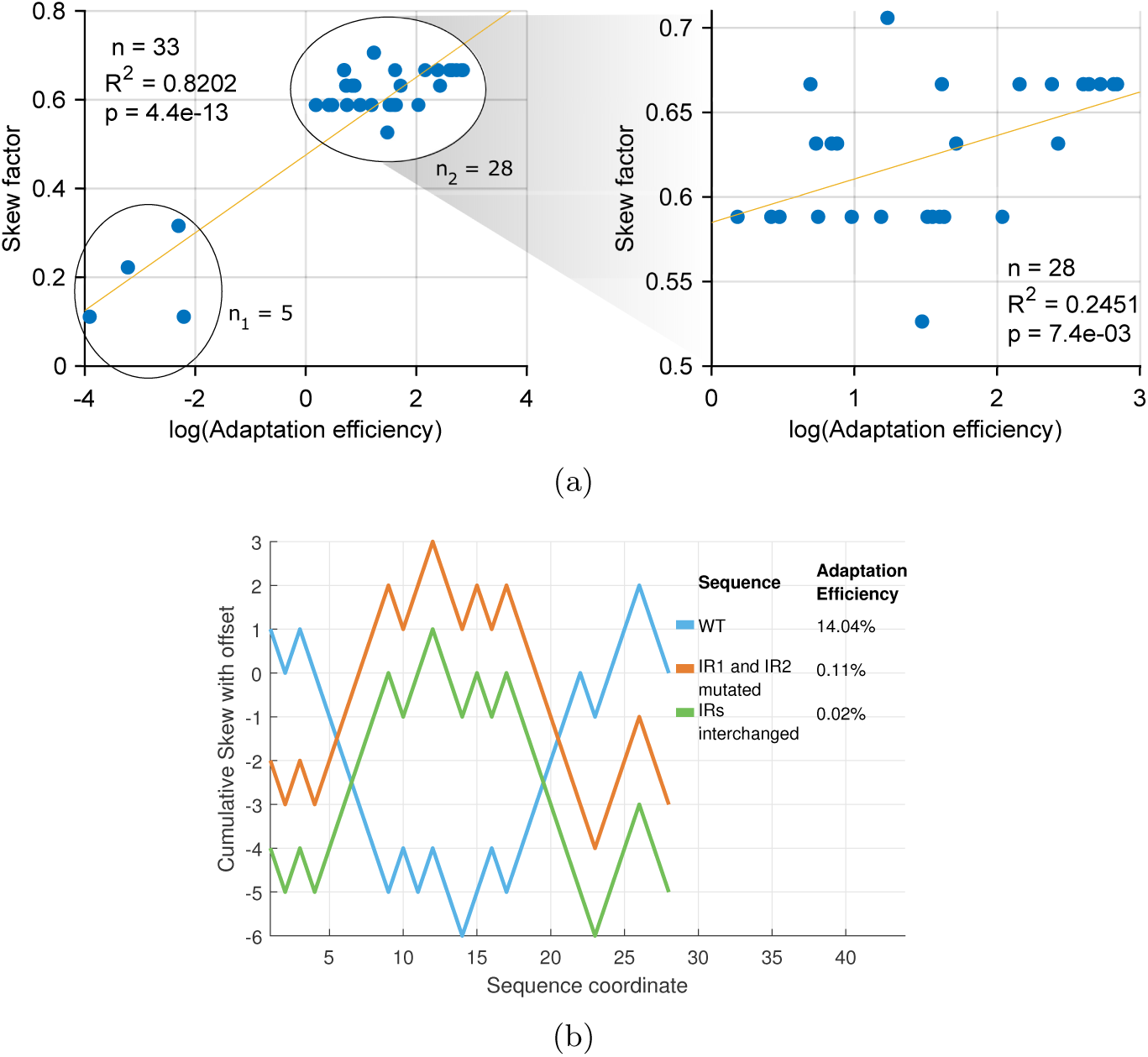
The figure illustrates the relationship between the adaptation efficiency and nucleotide arrangement. **(a)** The figure shows a positive correlation between the skew factor of the direct repeat and the logarithm of the adaptation efficiency of the CRISPR array (data from Ref. [58]), indicating that arrays with lower skew factors exhibit reduced adaptation efficiency and those with higher skew factors display higher adaptation efficiency. The data form two distinct clusters, one containing 5 points and the other 28 points; however, a positive correlation is still observed within the cluster (shown in the side panel). **(b)** The figure illustrates cumulative skew profiles for the RY nucleotide arrangement of the wild-type CRISPR array (blue), a mutant with substitution mutations in both inverted repeats (orange), and another mutant with the two inverted repeats interchanged (green). Their corresponding adaptation efficiencies are indicated in the legend.

The observed relationship (Fig. 10a) is further analyzed through cumulative RY skew profiles for mechanistic visualization. Figure 10b compares the cumulative RY-skew profiles and corresponding adaptation efficiencies for three CRISPR array variants: the wild-type sequence (5*^′^*-GTGT**TCCCCGC**GCCA**GCGGGGA**TAAACC-3*^′^* in blue curve), a mutant in which both inverted repeats are substituted (5*^′^*-GTGT**GAAAATA**GCCA**TATTTT C**TAAACC-3*^′^* in orange curve), and a variant in which the two inverted repeats are interchanged (5*^′^*-GTGT**GCGGGGA**GCCA**TCCCCGC**TAAACC-3*^′^* in green curve). The two inverted-repeats are 5*^′^*-TCCCCGC-3*^′^*(IR1), which is pyrimidine-enriched, and 5*^′^*-GCGGGGA-3*^′^*(IR2), which is purine-enriched.

The cumulative skew profile of the wild-type array shows a clear excess of pyrimidines across the left half (5*^′^*) of the direct repeat and purines across the right half (3*^′^*), which is similar to our canonical 5*^′^*-pyrimidines–purines-3*^′^*nucleotide arrangement. Within the sequence-dependent asymmetric cooperativity model, this asymmetry reduces the kinetic barrier heights near the center of the repeat around the 14*^th^* and 15*^th^* base pairs (Fig. 10b), creating a kinetically weak region that promotes unzipping initiation and bidirectional propagation. This kinetic favorability likely underlies the high adaptation efficiency observed for the wild-type array (14.04%). On the other hand, both the substituted-inverted-repeat and interchanged-inverted-repeat variants flipped the nucleotide motif from a 5*^′^*-pyrimidines–purines-3*^′^* arrangement to a 5*^′^*-purines–pyrimidines-3*^′^* configuration. This conversion makes the direct repeat less favorable for unzipping initiation, thereby hindering spacer integration and sharply reducing adaptation efficiency to 0.11% in the substituted inverted repeats variant and 0.02% in the interchanged inverted repeats variant.

Therefore, the severe decrease in adaptation efficiency due to the interchange of inverted repeats within the direct repeat argues against the necessity of secondary structures for spacer acquisition. Even after the interchange, cruciform-like secondary structures can still form; however, the directional RY-asymmetric orientation is disrupted, which markedly lowers the unzipping potential. Furthermore, several CRISPR repeat families, particularly many archaeal repeats, exhibit minimal or no palindromic nature and therefore are unable to form stable hairpin structures [59]. This argument demonstrates the necessity of the local nucleotide arrangement that favors unzipping over secondary-structure formation.

## 6 Discussion

In this study, we propose a mechanistic explanation for the integration of new spacers into CRISPR arrays by highlighting the functional roles of the leader–repeat junction, the first direct repeat, and the first repeat–spacer junction in determining both integration site specificity and adaptation efficiency. Our sequence analyses show that direct repeats are not compositionally random; instead, they exhibit a conserved 5*^′^*-pyrimidines to 3*^′^*-purines asymmetry, which confers high unzipping potential (Fig. S2). This directional asymmetric motif generates a characteristic kinetic landscape in which the central region of the repeat is most susceptible to unzipping initiation, whereas both ends form high–kinetic-barrier zones that impede further unzipping (Fig. 6 and 7). The resulting stalled intermediate generates an unstable denaturation bubble (step 2 in Fig. 5), which induces single-strand breaks at the 5*^′^*-ends of the bubble [51, 52, 53]. These asymmetric single-strand breaks provide room for the new spacer to integrate.

The implications of this mechanism are manifold. First, it rationalizes why the leader-repeat junction is universally preferred as the initial integration site, even in organisms lacking known guiding host factors such as IHF (Integration Host Factor) [22, 21]. In our model, purine enrichment at the 3*^′^*end of the leader and pyrimidine enrichment at the 5*^′^* end of the first repeat jointly create a high-kinetic barrier region that creates a kinetic bottleneck for the unzipping propagation and dictates the location for the spacer integration at the leader-repeat junction. Second, our results indicate that adaptation efficiency is not determined solely by Cas1–Cas2 compatibility, but is strongly modulated by the kinetic profile encoded in the direct repeat sequence itself. CRISPR arrays with direct repeats of higher skew factors (stronger directional *Y* → *R* asymmetry), reflecting higher unzipping potential, tend to exhibit higher adaptation efficiencies (Fig. 10 and Fig. 9).

Furthermore, our reinterpretation of prior mutagenesis studies provides experimental support for the proposed kinetic mechanism [29, 20, 58]. In these systems, perturbations that disrupt either the purine-rich or pyrimidine-rich half of the direct repeat lead to a pronounced loss of adaptation efficiency (see Fig. 10b). Importantly, we show that the sequence identity and the RY distribution pattern are predictive of functional outcomes, a feature overlooked in traditional sequence alignment-based analyses.

In conclusion, our study establishes a mechanistic model in which sequence-dependent DNA kinetics are used to explain both the site specificity and the variability in spacer integration efficiency. The results reveal the importance of the CRISPR array sequence architecture for integration. The conserved pyrimidine–purine asymmetry observed in CRISPR direct repeats suggests that these sequences are shaped by evolutionary constraints to maintain the asymmetric skew profile, extending beyond their established role in crRNA maturation. More broadly, these findings indicate that rationally tuning nucleotide asymmetry within the leader–repeat–spacer sequence could provide a strategy to engineer CRISPR arrays with controllable adaptation efficiencies, offering a potential route to optimize synthetic microbial immunity.

## Data Availability

The data, sequences, and algorithms used in the manuscript are available at: https://github.com/Sashik05/ Directional-purine-pyrimidine-asymmetry-in-CRISPR-repeats-facilitates-spacer-acquisition. git.

## Supplementary Information

### 1 Flow chart for skew factor

**Supplementary Figure S1:**
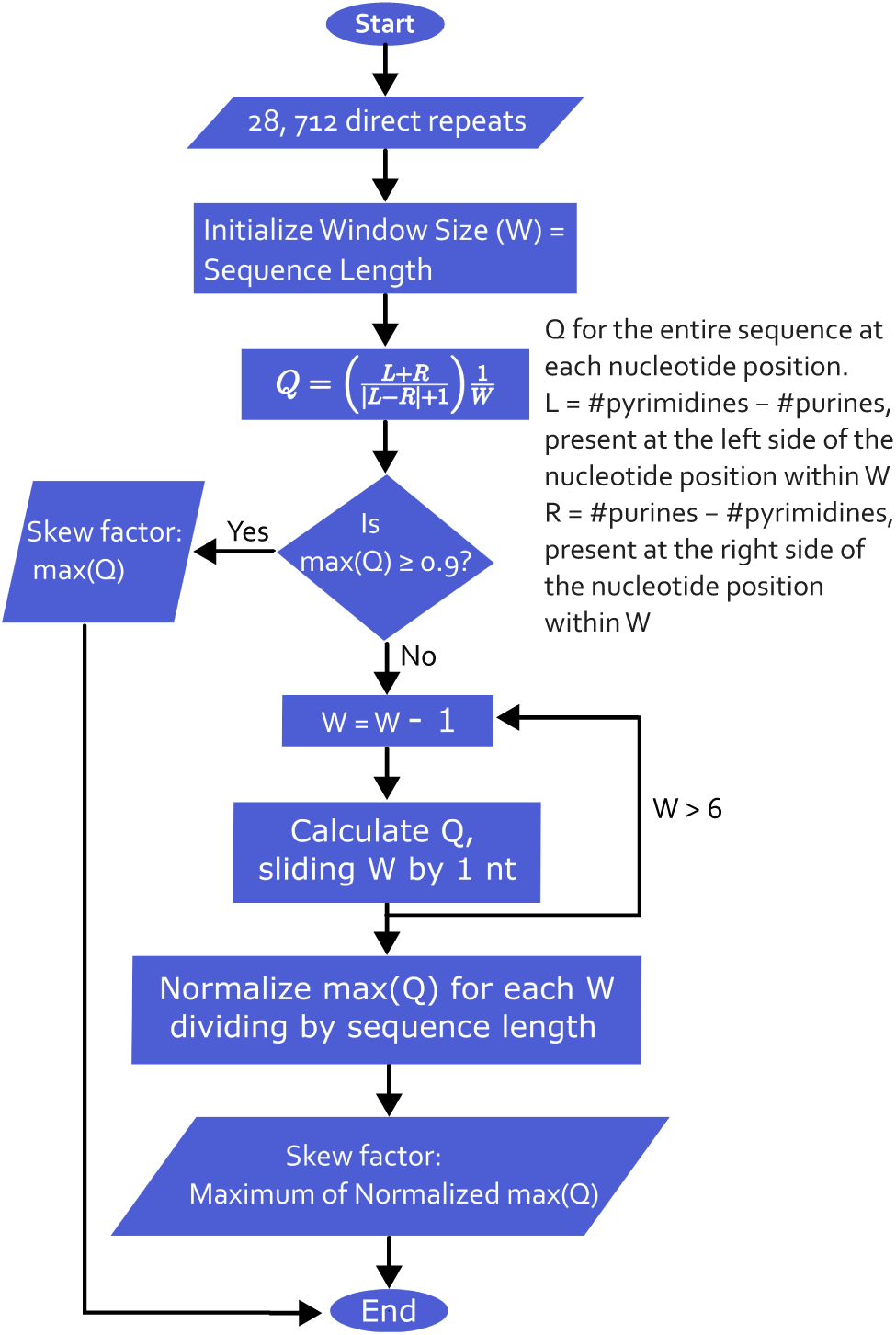
*The flow chart provides the algorithm to calculate* ‘skew factor’.

### 2 Parameter regime

The dynamics of unzipping is represented using a continuous-time Markov chain characterized by four parameters: the base pair formation rate (*p*), the dissociation rate (*r*), and two asymmetric cooperativity factors, catalytic (*α*) and inhibitory (*β*). Since the formation and dissociation of base pairs occur as intramolecular processes within the DNA duplex, these rates are assumed to be independent of strand concentration. The thermodynamic equilibrium constant of base pairing is expressed as

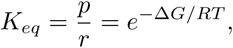

where Δ*G* is the Gibbs free-energy difference between the bonded and unbonded states, *R* is the universal gas constant, and *T* is the absolute temperature. Sequence-specific thermodynamic parameters are obtained from experimentally determined nearest-neighbor free energies reported by SantaLucia *et al.* [60]. The thermodynamic parameters, *p* and *r* are considered to be 5*k* and 0.66*k*, respectively, where *k* is a scaling factor of the order of 10^6^ s*^−^*^1^ [61]. These thermodynamic parameters are chosen at 340 K (*T_m_* − 33*K*). *T_m_* is the average melting temperature of any base pair, *T_m_* = 373.22*K*. The asymmetric cooperativity parameters are introduced as multiplicative factors modifying the transition rates of neighboring base pairs, where *α* (*α* = 5) represents catalytic enhancement and *β* (*β* = 0.2) represents inhibitory interaction. The obtained results are consistent with a wide range of parameter values as long as they satisfy α ≠ β to introduce directional asymmetry in the unzipping kinetics.

### 3 Fastest and slowest sequences

Mean First Passage Time (MFPT) is defined as the expected time required for a Markov chain (*X_t_*)*_t≥_*_0_ to reach a specified target state within our state space *S*, defined in ‘Unzipping pathway’. In this study, the target state corresponds to the fully unzipped DNA configuration, while the initial state represents the fully zipped double-stranded DNA. MFPT is computed by inverting a modified generator matrix Q*^′^*, obtained by removing the row(s) and column(s) corresponding to the target state, as expressed in the following equation [39].

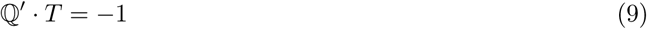

The resulting residence time matrix **T** has elements *t_ij_*, which represent the expected time spent in state *j* when the Markov chain starts from state *i* during its progression toward the target state. The MFPT of the transition to the target state from state *i* can be obtained by adding the residence times of the transient states.

**Supplementary Figure S2:**
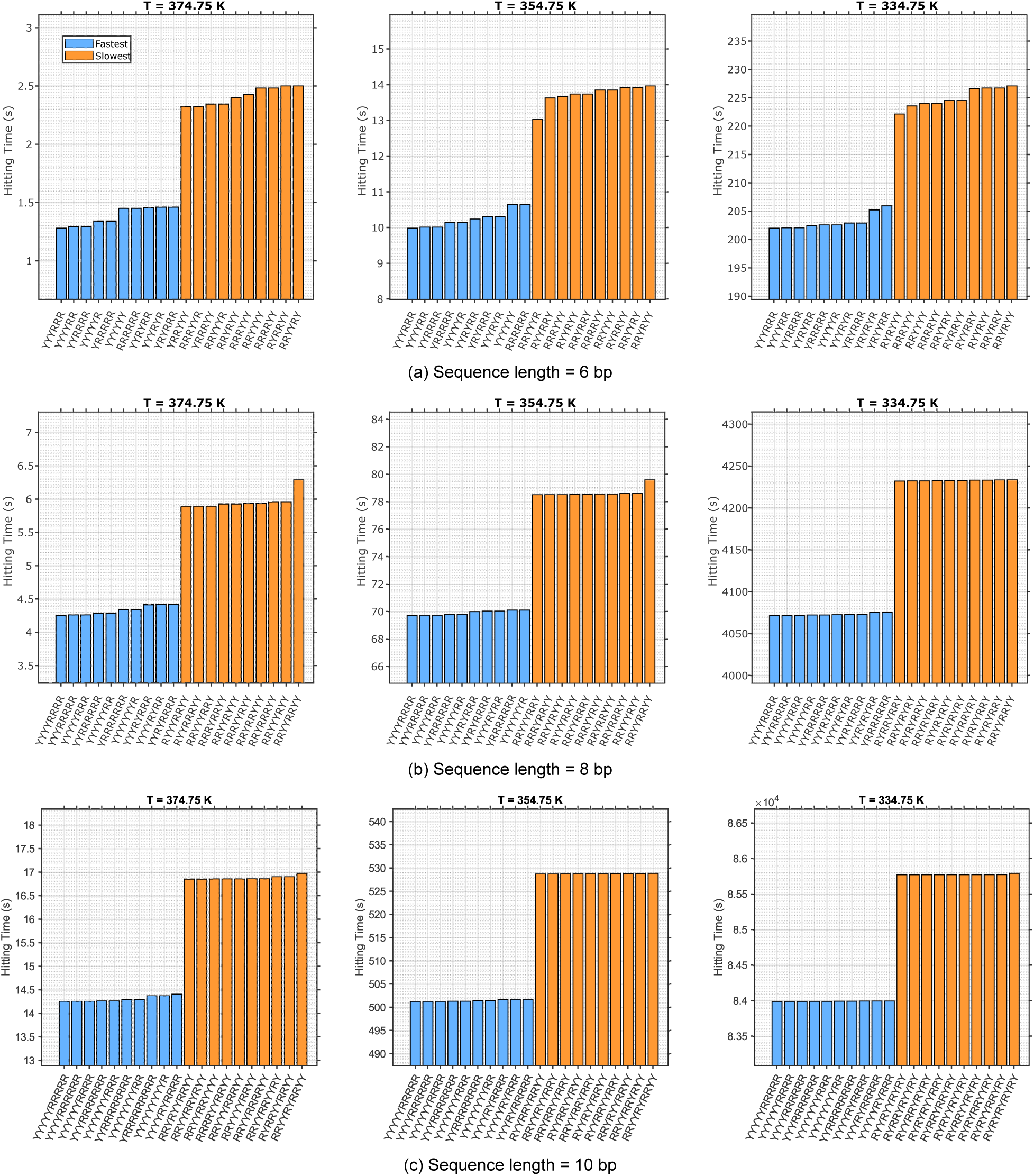
The figure illustrates the mean first passage time (MFPT) for unzipping, time taken for separation of two ssDNA strands from a dsDNA, of the fastest (blue bars) and slowest sequence (orange bars) of sequence lengths 6, 8 and 10 bp at different temperatures varying from 374.8 K (near average melting temperature considering both G-C and A-T) to 334.75 K in steps of 20 K. All the fastest sequences feature pyrimidines in 5^′^-half and purines in 3^′^-half. The slowest ones have alternating stretches of purines and pyrimidines, starting with purines from the 5^′^-end.

### 4 Relationship between skew factor and unzipping potential

Figure S3 illustrates the relationship between the skew factor, which dictates the sequence motif, and the rate of unzipping, which reflects the functional characteristics of the sequence. The figure indicates an approximately linear relationship between the skew factor and the unzipping potential. However, degeneracy in the skew factor is observed for sequences with low values of the skew factor.

**Supplementary Figure S3:**
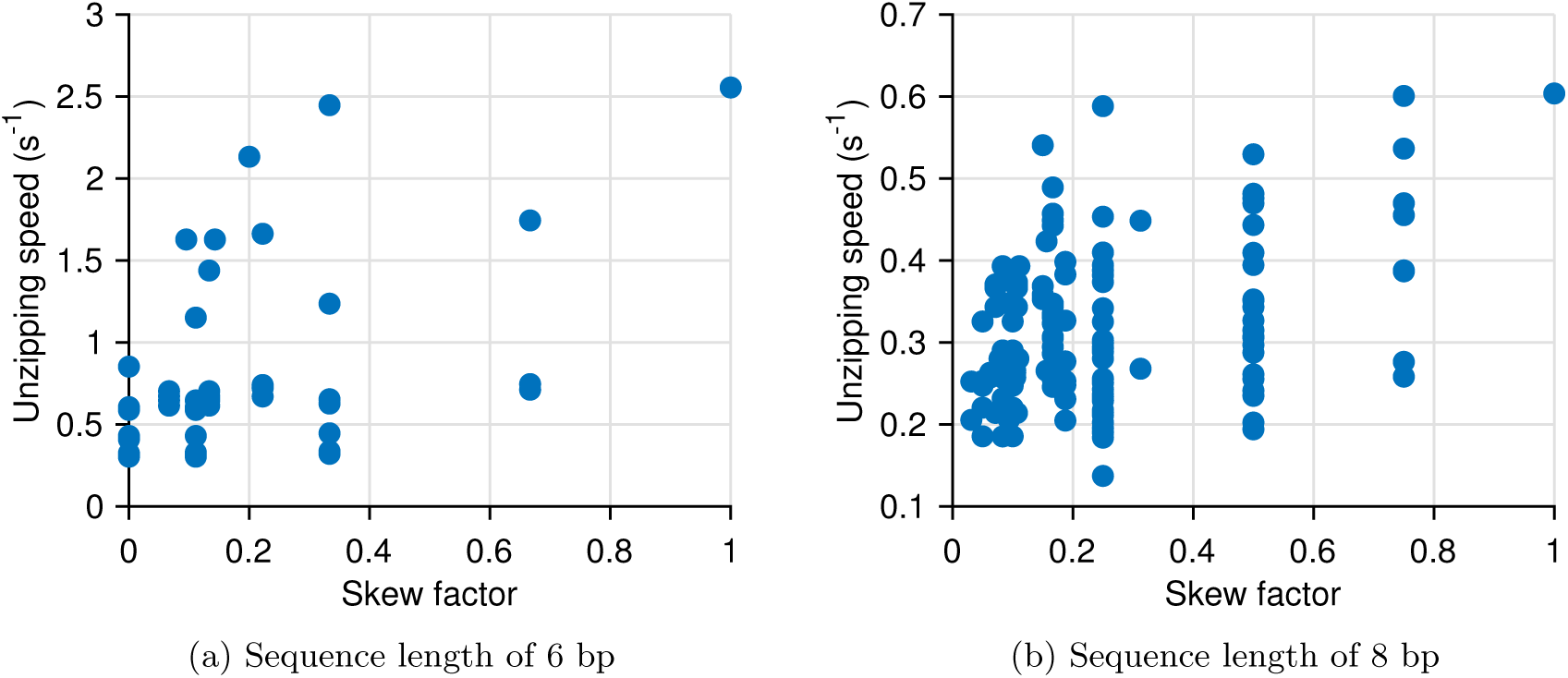
The figure illustrates the relationship between the skew factor and the unzipping speed, defined as the reciprocal of the mean first passage time (MFPT) of the unzipping process. In the present analysis, however, the thermodynamic parameters corresponding to T = 350 K are used, with α = 5 and β = 0.2, consistent with the parameter values used in the calculation of the fastest and slowest sequences. **(a)** for all possible sequences of sequence length 6 bp. **(b)** for all possible sequences of sequence length 8 bp.

### 5 Demonstration of unzipping dynamics

**Supplementary Figure S4:**
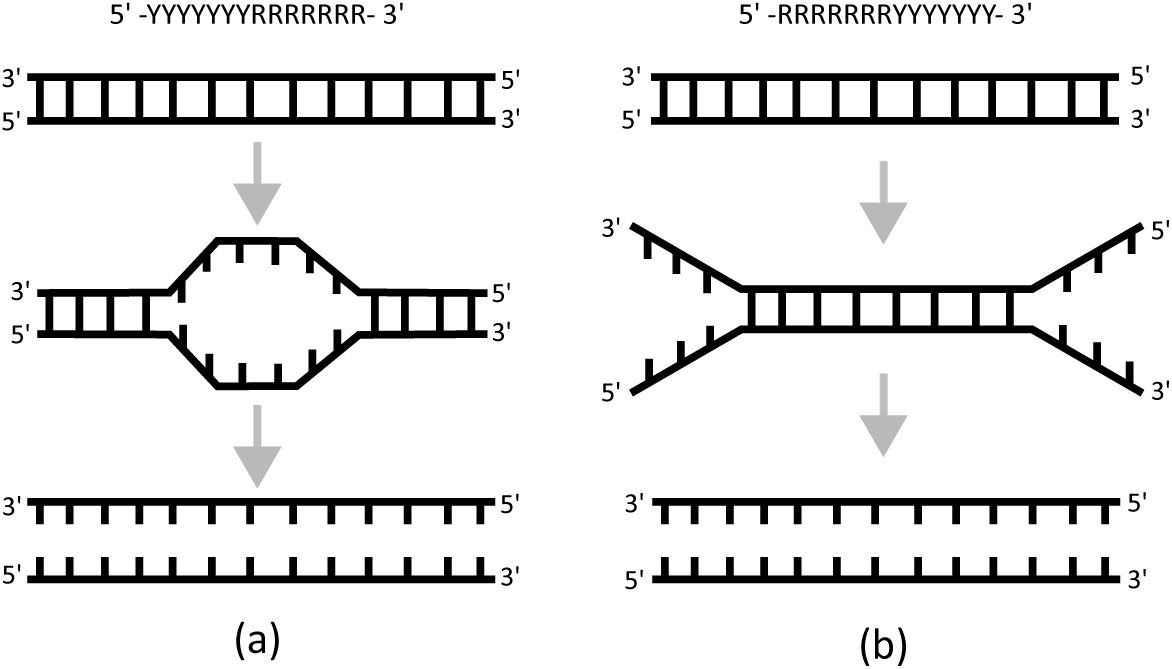
Mechanistic illustration of the unzipping process for two extreme sequences defined by the skew factor. (a) Unzipping of 5^′^−YYYYYYY RRRRRRR−3^′^ (skew factor = 1). The process initiates at the middle base pairs, which possess lower kinetic barrier heights (see Methods), leading to the formation of a central bubble. The bubble subsequently expands bidirectionally toward both ends, ultimately yielding two separate single strands. (b) Unzipping of 5^′^ − RRRRRRRYYYYYYY − 3^′^ (skew factor = 0). In this case, the middle base pairs exhibit higher kinetic barrier heights that hinder dissociation. As a result, unzipping proceeds from the ends, forming two Y-shaped forks. This configuration is unstable and may eventually lead to a double-strand break.

### 6 Detailed analysis of Fig. 9

In Fig. 9a, the mutant types WT, G1A C30T, G1T C30A, G1C C30G, G1A, G1T, G1C, C30A, C30T, C30G, B4, B7, B9, B6.1, Insertion and Deletion exhibit broadly similar cumulative RY-skew profiles. In all of these cases, a pronounced V-shaped feature is observed near sequence coordinate ∼15, accompanied by Λ-like and zig-zag regions spanning approximately coordinates 3–6 and 40–53. Some of these mutants, like WT, G1C C30G, G1A, G1C, C30A, C30T, C30G, have nearly the same reads of insertion events with a prominent leader–repeat integration site near coordinate ∼11 and a repeat–spacer integration site near coordinate ∼40. These mutants also display low insertion events around coordinates ∼6 and ∼46. However, the mutants G1T C30A, G1T, B4, B7, B9, and B6.1 don’t have appropriate insertion hotspots, despite exhibiting a significant cumulative RY skew profile.

Deviations from this characteristic skew profile, B1 and B2 (Fig. 9a) show strong purine enrichment across the array within the direct repeat, indicating a low propensity for unzipping initiation at this site. Although a shallow V-shaped profile is present within the spacer region and may locally initiate unzipping, further propagation toward the 5*^′^* end is disfavored because the upstream (5*^′^*) sequence is purine-rich and therefore dominated by the right-asymmetric mode of cooperativity. Therefore, the arrays (B1 and B2) don’t have any proper insertion sites. The B6, B6.2, and B6b mutant types exhibit multiple shallow V-shaped local profiles that may facilitate unzipping initiation at several locations, but they lack robust stall sites, resulting in the absence of distinct, high-read integration hotspots. However, they exhibit multiple insertion sites with low read counts.

Furthermore, B8, B10, and Non-CRISPR (in Fig. 9a) display similar overall cumulative RY-skew profiles. However, B10 does not show any high read insertion sites. Short WT shows a V-shape skew profile and insertion sites at the both ends of the mutant array. While short B6 doesn’t have any proper unzipping initiation sites due to the absence of V-shape high skew profile. The Non-CRISPR-mut array exhibits pyrimidine enrichment up to approximately coordinate 50 and a V-shaped skew profile within the spacer region, which can favor unzipping initiation and propagation toward the 5*^′^*end. However, this array lacks stall sites along the unzipping direction, resulting in the absence of well-defined insertion sites.

A consistent sequence dependence is also observed in another experimental study, shown in Fig. 9b. Most mutant arrays show a characteristic double V-shaped cumulative RY skew profile near the center of the direct repeat, which facilitates efficient unzipping initiation. In addition, a pronounced Λ-shaped feature near sequence coordinate ∼11 and a zig-zag region spanning coordinates ∼38–46 act as kinetic pause sites and coincide with dominant insertion hotspots. Following this skew profile, the leader–repeat integration site is observed near the 11*^th^* position, while the repeat–spacer integration site occurs around coordinates 44–46.

B1 and B2 in Fig. 9b exhibit a V-shaped skew profile at the 11*^th^* position rather than a Λ-shape (pause site signature), which results in the absence of prominent insertion sites. Moreover, the presence of an additional V-shaped feature within the spacer region can reshape the overall unzipping landscape and, in turn, influence insertion sites. As a result, repeat-spacer insertion sites in these mutants are distributed across multiple nearby coordinates (e.g., 45, 46, 47, and 54), corresponding to positions flanking the spacer-associated V-shaped skew profile. Therefore, these observations show that the insertion sites are not strictly confined to the leader–repeat and repeat–spacer junctions but are influenced by the local sequence composition.

### 7 RY asymmetric signatures for integration events outside CRISPR biology

Similar RY-asymmetric signatures are also observed near integration sites outside CRISPR biology, including those associated with the Tn7 transposon, Class C attC Group II introns, integron cassettes, and retroviral integration [62, 63, 64, 65, 66, 67], suggesting a common structural or sequence-based mechanism similar to our proposed model. A few well-characterized integration sites reported in Refs. [63, 68, 69], the attTn7 sites shown in Fig. S5a, represent highly conserved insertion hotspots for the Tn7 transposon, whereas the attC site shown in Fig. S5b enables the integration of gene cassettes into integrons. These attC sites are also capable of forming hairpin-like structures [67]. We also identify an RY-asymmetric nucleotide distribution near retroviral integration sites, including HIV-1, from previously reported sequence data [70, 71], shown in Fig. S6. These observations show the requirement of the RY asymmetric distribution of the target site for integration.

**Supplementary Figure S5:**
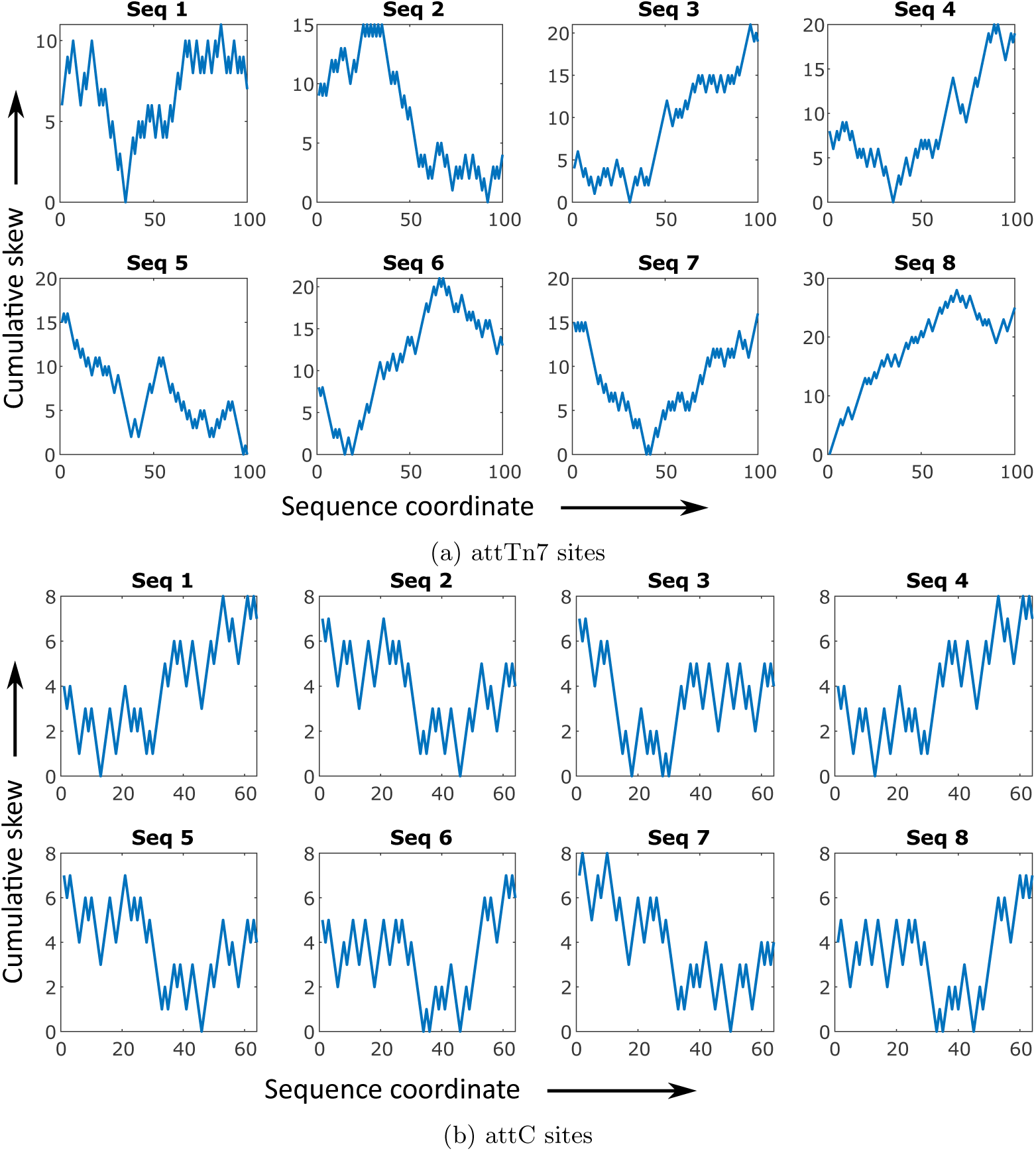
Cumulative RY-skew profiles of experimentally characterized integration sites outside CRISPR arrays. Despite their distinct biological contexts, these sequences exhibit cumulative RY-skew patterns that resemble those observed in the leader–first repeat–first spacer region of CRISPR arrays. Most profiles display a characteristic V-shaped structure, and shifts or distortions in this V-shape are associated with changes in the location and efficiency of integration sites, although a quantitative relationship was not reported in the original studies. (a) Cumulative RY-skew plots for eight attTn7 sites, which serve as conserved insertion sites for Tn7 transposons. (b) Cumulative RY-skew plots for eight attC sites that mediate the integration of gene cassettes into integrons; the sequences shown correspond to different mutant variants.

**Supplementary Figure S6:**
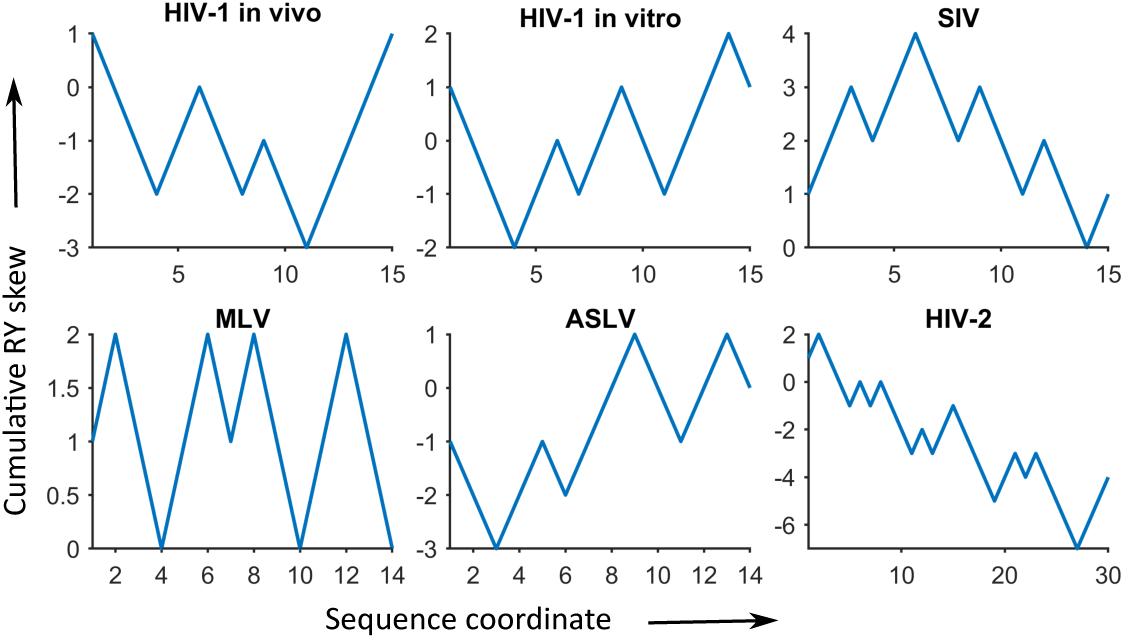
Illustration of RY cumulative skew profile of the integration sites for the retroviral elements, such as HIV (Human immunodeficiency viruses), SIV (Simian Immunodeficiency Virus), MLV (Murine Leukemia Virus) and ASLV (Avian Sarcoma and Leukosis Virus).

